# An open source 16-channel fluidics system for automating sequential fluorescent *in situ* hybridization (FISH)-based imaging

**DOI:** 10.1101/2022.03.23.485524

**Authors:** Zhaojie Deng, Brian J. Beliveau

## Abstract

Fluorescent *in situ* hybridization (FISH) can provide spatial information about DNA/RNA targets in fixed cells and tissues. However, the workflows of multiplexed FISH-based imaging that use sequential rounds of hybridization quickly become laborious as the number of rounds increases because of liquid handling demands. Here, we present an open-source and low-cost fluidics system that is purpose built for automating the workflows of sequential FISH-based imaging. Our system features a fluidics module with 16 addressable channels in which flow is positive pressure-driven and switched on/off by solenoid valves in order to transfer FISH reagents to the sample. Our system also includes a controller with a main printed circuit board that can control up to 120 solenoid valves and allows users to control the fluidics module via serial communication. We demonstrate the automatic and robust fluid exchange with this system by targeting the alpha satellite repeat in HeLa cell with 14 rounds of sequential hybridization and imaging. We anticipate that this simple and flexible system will be of utility to researchers performing multiplexed *in situ* assays in a range of experimental systems.

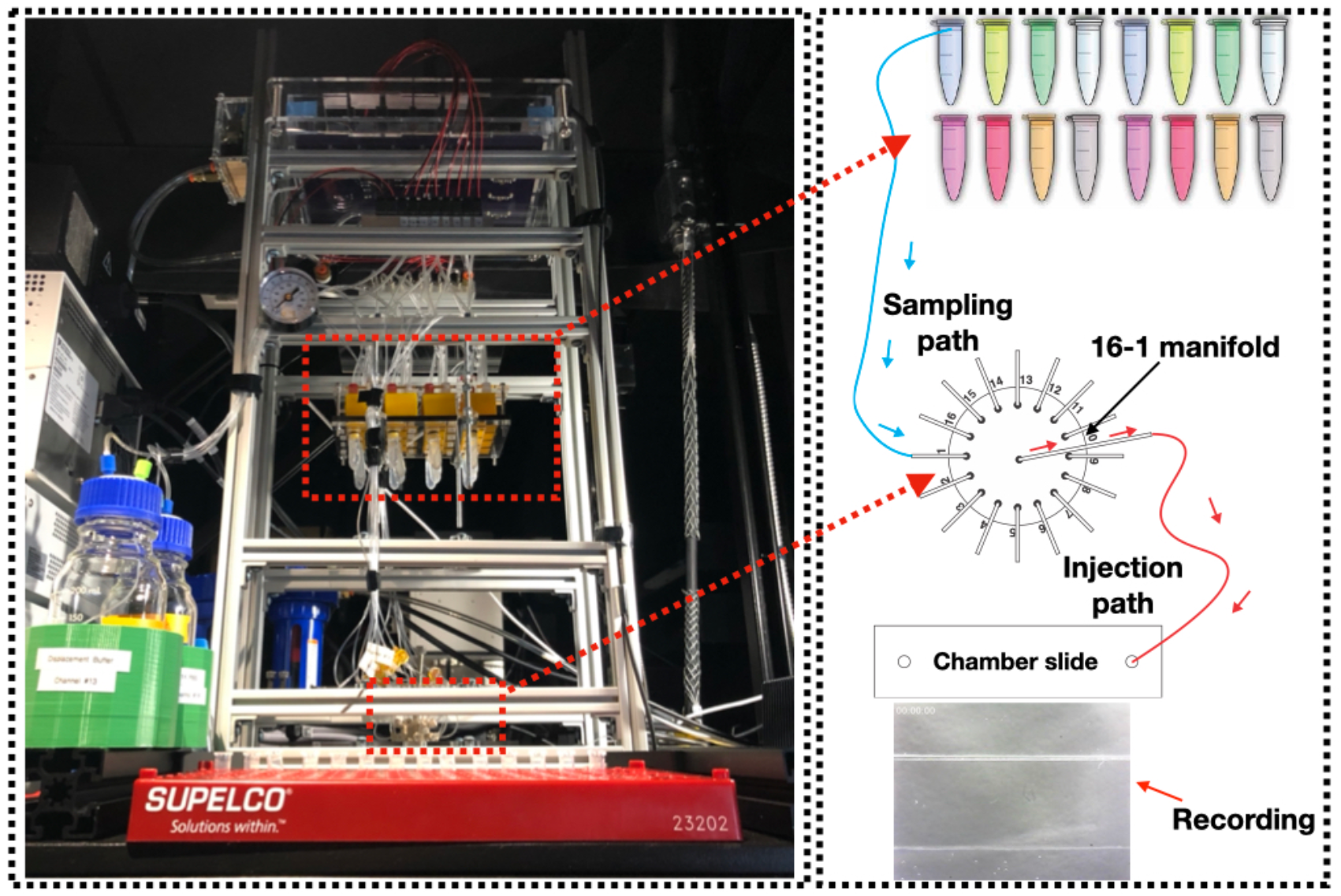

**Specifications table:** 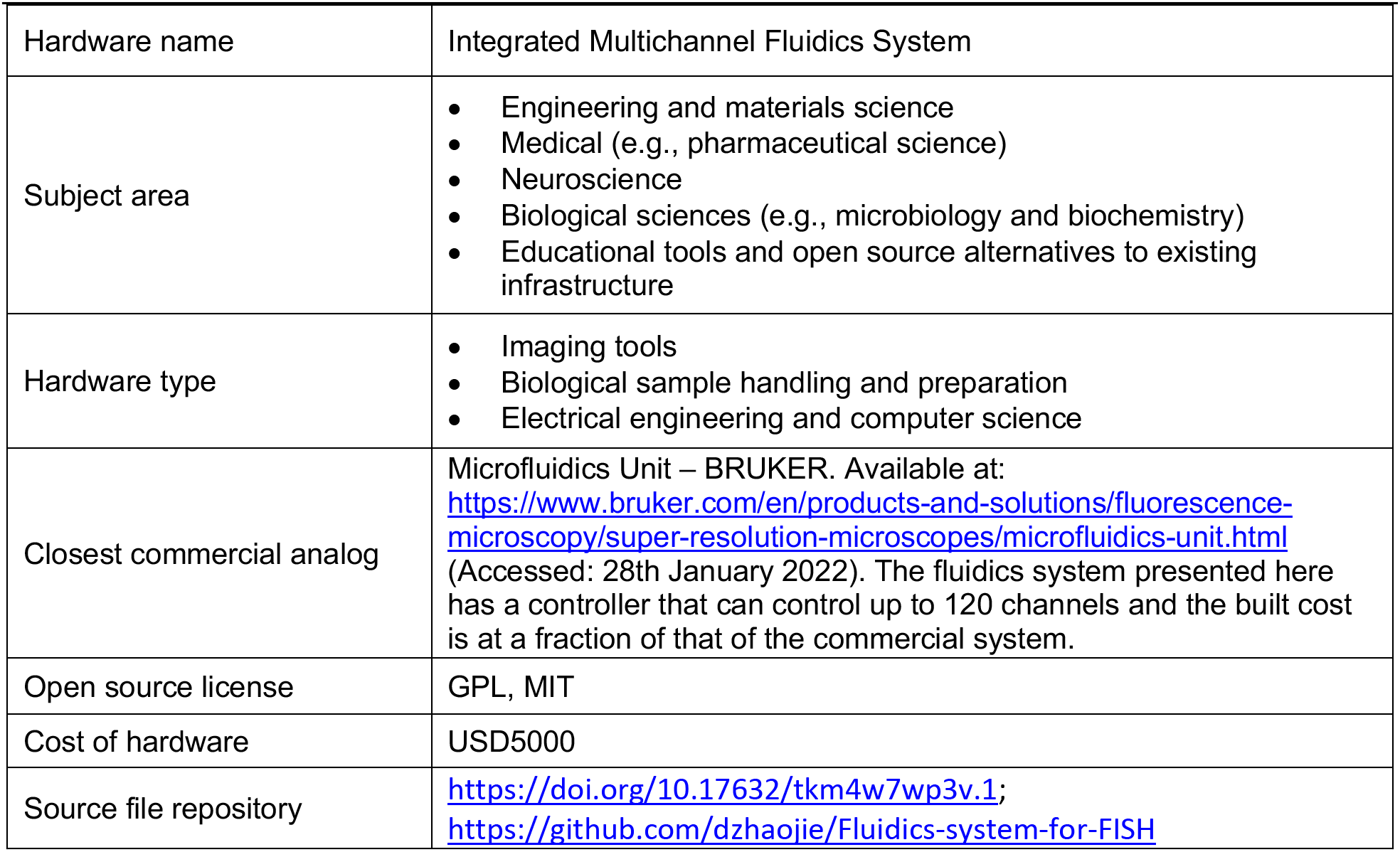

## 1. Hardware in context

Fluorescent *in situ* hybridization (FISH) is a technique that uses fluorescently labeled nucleic acid probes to target genomic and/or transcriptomic sequences of interest in cells and tissues through hybridization. After hybridization, the locations and abundances of the targets can be detected by fluorescence microscopy. Therefore, FISH enables researchers to obtain spatial genomic and transcriptomic information relevant to their research. Recent developments in FISH technology have focused on improving its multiplexing capability in order to efficiently detect more than a dozen genes and/or RNAs in one experiment by either spectral encoding [1], [2] or sequential hybridization [3], [4], [5], [6]. Recent advances in multiplexing strategies involve combining spectral encoding of the DNA/RNA targets with sequential hybridization of FISH readout probes to detect thousands of DNA/RNA species [7], [8], [9], [10]. Sequential FISH procedures usually involve many rounds of hybridization. During each round, readout probes are hybridized and imaged and their signal is removed by either one or two of these methods: photobleaching, toehold displacement of the readout probes, chemically stripping of the readout probes. Protocols for sequential FISH-based imaging are usually complicated and involve multiple liquid handling and/or exchange steps in each hybridization round. Therefore, they are time consuming and labor-intensive for manual implementation, preventing them from being widely adopted.

Commercially available liquid handling solutions [11], [12], [13] are either hard to adapt to sequential FISH-based imaging or are expensive when the rounds of imaging need to be scaled up. An open-source integrated automation solution capable of handling more than a dozen rounds of fluid exchange for sequential hybridization would help address this challenge. Integrated microfluidics platforms have been demonstrated to perform all assay steps in an entire FISH protocol [14]. However, these platforms can only perform up to a dozen hybridization rounds and they are difficult to scale up. Other solutions utilize peristaltic pumps combined with rotative valves to automate fluid exchanges up to 24 rounds in imaging chambers [15] or with Lego-hardware to support up to 128 syringe pumps to input reagents into a peristaltic pump driven flow system [16]. While these solutions provide high-throughput automation, peristaltic pumps offer less stability in flow rate control for long-term experiment like sequential FISH more than a dozen rounds [17]. On the other hand, pressure-driven provides high stability and pulseless flow [17]. Here, we present an open source fluidics system with 16 channels individually driven by positive pressure for automation of sequential FISH-based imaging protocols. We show robust fluid exchange with the system through 14 rounds of fluorescent oligo hybridization (2 other channels were used for washing buffer and probe displacement buffer) to readout the primary probes at the alpha satellite in Hela cells. While maintaining a comparable dead volume (100 μL) to commercial ones, this open source standalone fluidics system offers a custom designed sample loading module (no extra tool required) for conveniently loading probe solutions to significantly reduce experiment prep time. It can also be integrated with a microscope enabling users to run/test FISH-based imaging protocols easily in their own laboratories.

## 2. Hardware description

### 2.1 Design Overview

The fluidics system here was designed with full automation and minimized reagent cost in mind. It also was built with low cost and widely available components. It operates via positive pressure-driven flow to introduce FISH reagents to the sample. The positive pressure can come from any common laboratory compressed air source. The whole system includes a fluidics module and a PCB controller. An overview of the fluidics module is shown in Fig. 1. A compressed gas line runs first through a filter module (component R2) and then a pressure regulator (component R3). The pressurized gas is then distributed into 16 individual lines by a splitting manifold (component R6) and 2 solenoid valve and manifold assemblies (component SV1&SV2). These 16 gas lines (T4) can be individually addressed to pressurize a designated reservoir filled with FISH probes or washing buffer. The 16 fluid lines (T2) coming out of the reservoirs are then combined by a mini manifold (with 16 inlets and 1 inlet) and become a single fluid line connected directly to the inlet of a sample chamber. A sample loading module (not shown in the diagram) composed of an array of 16 custom designed and 3D-printed Eppendorf tube adapters (component E1 here) was designed and assembled for easy sample loading using conventional 2 mL Eppendorf tube (component b). A structural frame (not shown in the diagram) was built to organize all the components into a standalone machine.

**Fig. 1.**
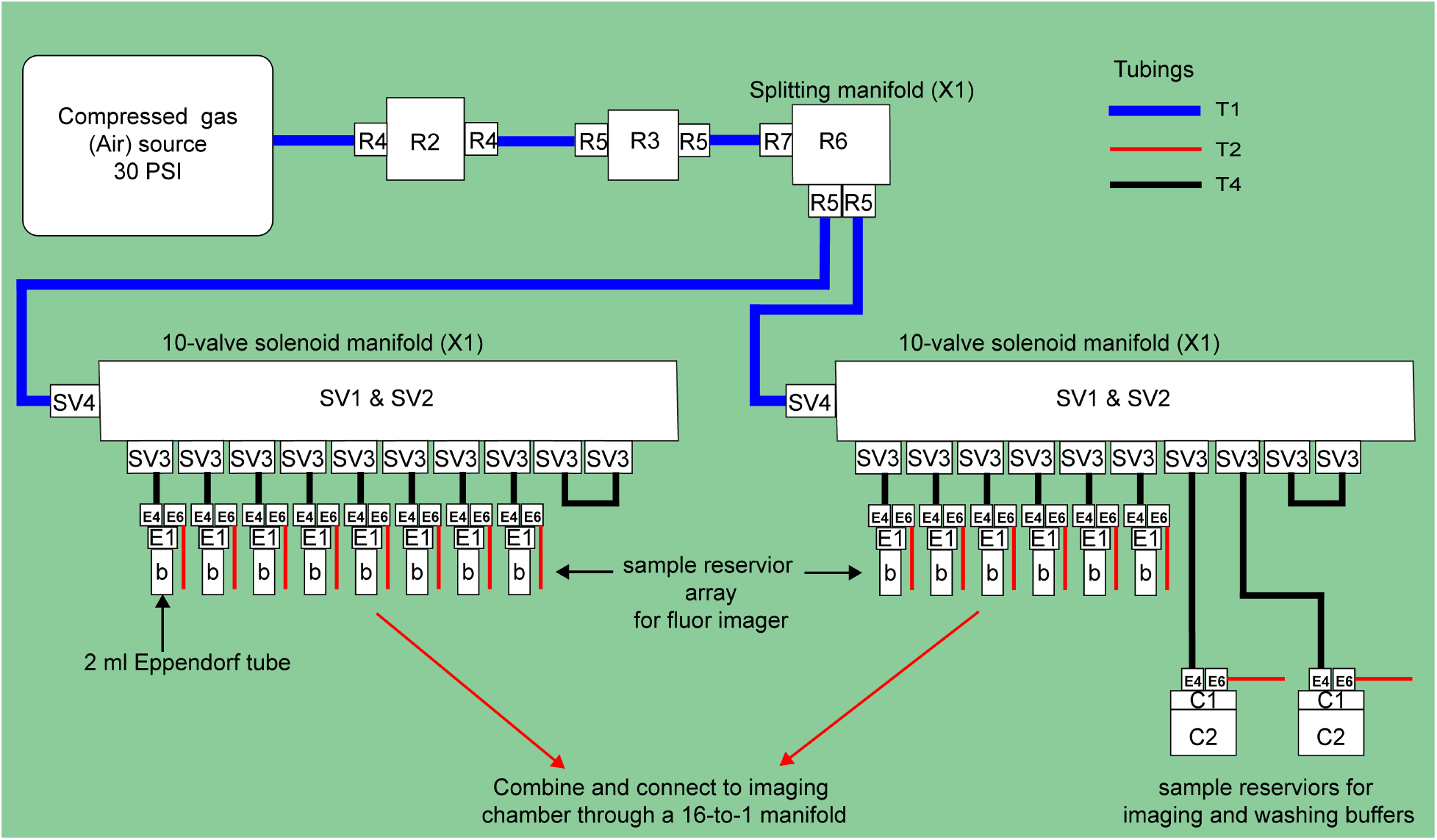
Overall schematic of the fluidics module (pneumatic and fluidics components) designated by letters and numbers (as referenced to the Bill of materials)

### 2.2 Controller PCB

A controller was designed for controlling the on/off of the solenoid valves, the pressure regulator, and a flow rate sensor, as well as handling serial communication with the microscope PC running the imaging routines. The main PCB of the controller is based on the design in the open-source microfluidics control system by Craig Watson [18]. It uses Espressif ESP32 devkit C as microcontroller and PCA9698 as I/O expander. Here, the PCB can address up to 120 solenoid valves after adding 2 more PCA9698 chips. Additionally, an interface is also added for a flow rate sensor (MFS, ElveFlow) powered by 9 V and it uses analog pins to read in the voltage output by the sensor. The 9 V is provided by a 12 V to 9 V regulator.

The interface for the pressure regulator is designed to control a commercial digital pressure regulator from (MPV, Proportion-Air). This controller handles all the messages from the PC through USB serial communication to operate the fluidics system.

### 2.3 User Interface

A user interface for flow rate calibration was developed using PyQt, Fig. 2. It has 4 tabs. The first 3 tabs are designed to run the main procedures for flow rate calibration including channel priming, flow rate calibration, and flow rate verification. The 4^th^ tab is for selecting an individual channel to inject a solution for a user-defined amount of time for illuminating bubbles before setting up an imaging experiment. Calibration data and the resulting injection time (from a user defined volume to inject) for each channel will be saved as *.npy* files by the app. In this demonstration with a use case in our lab, the resulting injection times for each channel in the *.npy* file named *WaitTime.npy* can be transferred into a NIS-Element (Nikon software for imaging) Macro file and used in NIS-Element’s ND acquisition with time-lapse imaging. Nikon NIS-Element macro codes are provided in the project’s repository for running the 14-round sequential FISH imaging on pan alpha satellite in HeLa cell as a demonstration experiment with the fluidics system.

**Fig. 2.**
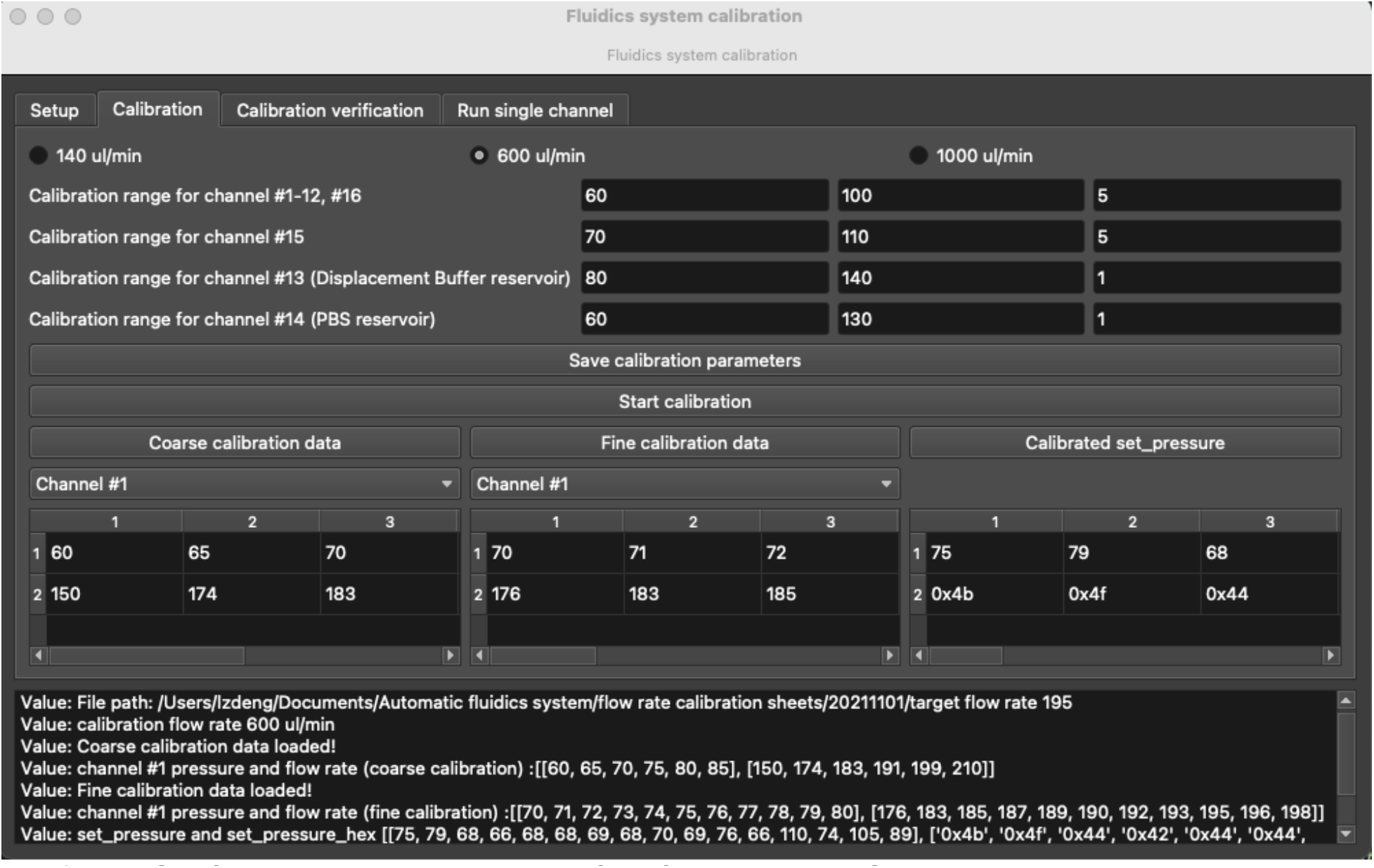
The GUI for the automatic calibration of the fluidics system. Shown here is the calibration tab with calibration parameters configured for target flow rate 600 μL/min.

## 3. Design files summary

| Design file name | File type | Open source license | Location of the file |
| --- | --- | --- | --- |
| bill_of_materials.xlsx | XLSX | GPL | Available with the article |
| fluidics_system_controller_PCB_design_files.zip | PCB files, KiCAD | GPL | https://doi.org/10.17632/tkm4w7wp3v.1; https://github.com/dzhaojie/Fluidics-system-for-FISH |
| Eppendorf_tube_adaptor_3D_print_files.zip | STL | GPL | https://doi.org/10.17632/tkm4w7wp3v.1; https://github.com/dzhaojie/Fluidics-system-for-FISH |
| laser_cut_acrylic_panel_organizer.ai | Illustrator file | GPL | https://doi.org/10.17632/tkm4w7wp3v.1; https://github.com/dzhaojie/Fluidics-system-for-FISH |
| controller_case_laser_cut.zip | 3dm file | GPL | https://doi.org/10.17632/tkm4w7wp3v.1; https://github.com/dzhaojie/Fluidics-system-for-FISH |
| case_for_pressure_regulator_laser_cut.pdf | PDF | GPL | https://doi.org/10.17632/tkm4w7wp3v.1; https://github.com/dzhaojie/Fluidics-system-for-FISH |
| large_reservoir_holder.stl | STL | GPL | https://doi.org/10.17632/tkm4w7wp3v.1; https://github.com/dzhaojie/Fluidics-system-for-FISH |
| user_interface_app_for_flow_rate_calibration.zip | Python script | MIT | https://doi.org/10.17632/tkm4w7wp3v.1; https://github.com/dzhaojie/Fluidics-system-for-FISH |
| NIS-Element Macro codes.zip | .mac | MIT | https://doi.org/10.17632/tkm4w7wp3v.1; https://github.com/dzhaojie/Fluidics-system-for-FISH |
| Fluidics System Controller with ESP32 | C++ code | MIT | https://doi.org/10.17632/tkm4w7wp3v.1; https://github.com/dzhaojie/Fluidics-system-for-FISH |

**bill_of_materials.xlsx**: There are 8 sheets in this file. Except the sheets named ‘Tubings’ and ‘Remaining components’, each sheet lists components for a module and/or assembly described in the step-by-step building guide in the supplementary materials.

**fluidics_system_controller_PCB_design_files.zip**: KiCAD design files for the controller PCB

**Eppendorf_tube_adaptor_3D_print_files.zip:** STL files for 3D printing the Eppendorf tube adaptor and the slot panel that is designed for arranging and securing the adapters

**laser_cut_acrylic_panel_organizer.ai**: Illustrator file for laser cutting the panels for assembling the structural frame and the Eppendorf tube adapter array

**controller_case_laser_cut.3dm**: Rhinoceros design file for laser cutting the controller case

**case_for_pressure_regulator_laser_cut.pdf**: pdf file for laser cutting the pressure regulator case

**large_reservoir_holder.stl**: STL file for 3D printing the large reservoir holders for securing large glass bottles for the washing buffer and displacement buffer used in the 14-round sequential FISH imaging demonstration experiment

**user_interface_app_for_flow_rate_calibration.zip**: Python scripts for the user interface app for automatic flow rate calibration

**NIS-Element_Macro_codes.zip**: Folder with NIS-Element Macro code for running the fluidics system and routines for the fluid exchange in the 14-round sequential FISH imaging demonstration experiment **Fluidics System Controller with ESP32:** Folder with **s**ource code for the firmware of the controller PCB

## 4. Bill of materials summary

A separate bill of materials file named “bill_of_materials.xlsx” is included in the supplementary materials with multiple sheets. Each sheet lists parts for an individual module and/or assembly that makes up the fluidic system. The table below lists all the parts from that file. The cost for the component is listed in currency USD.

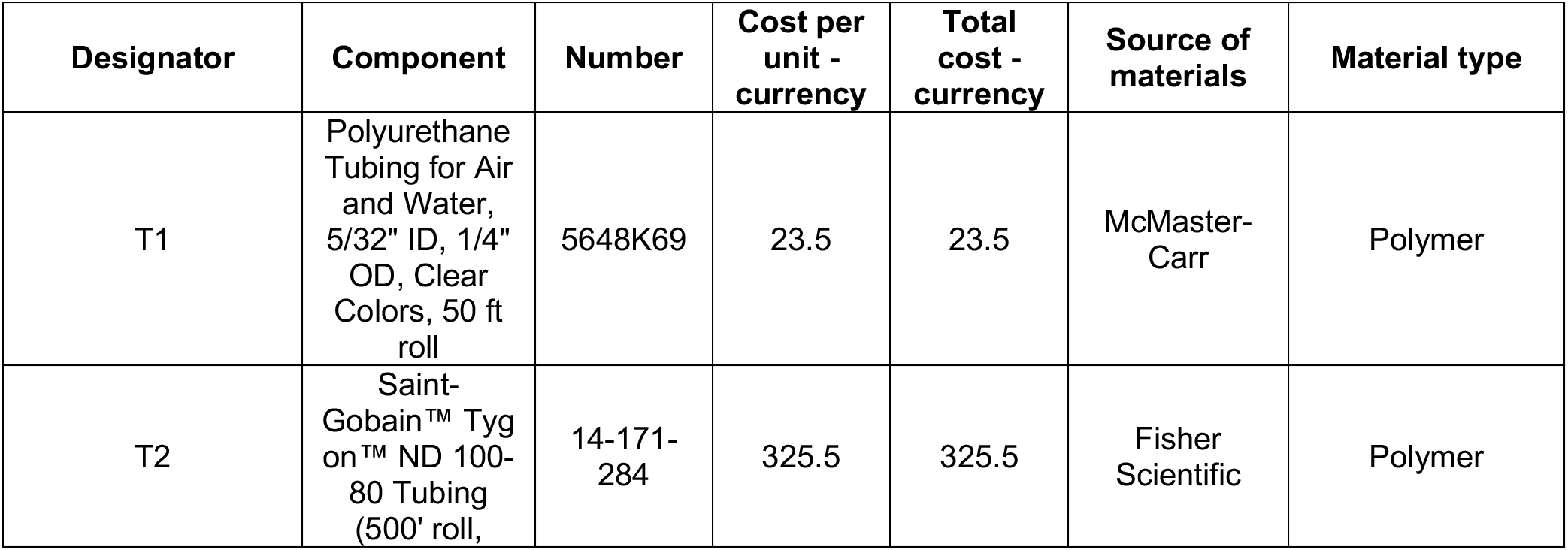
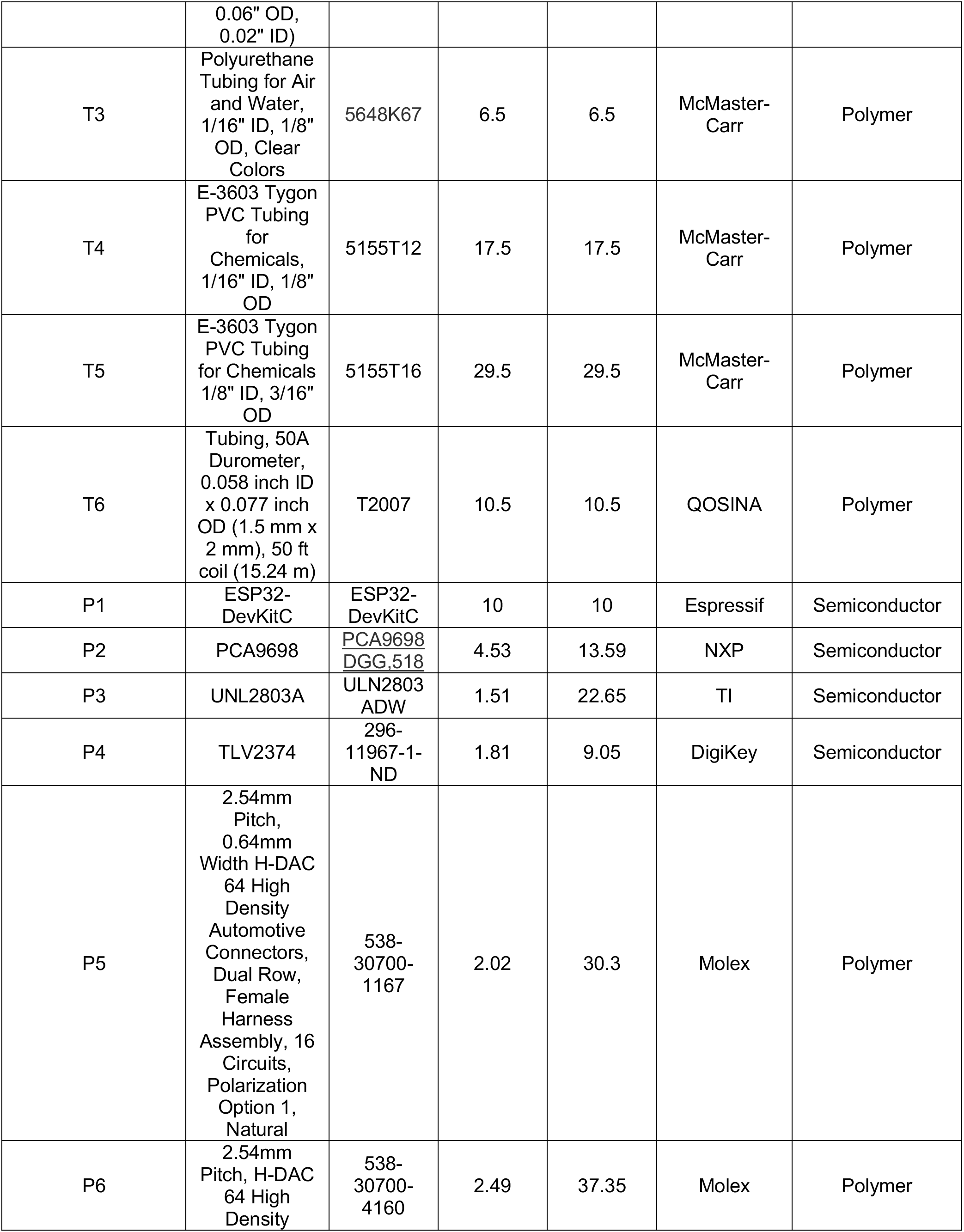
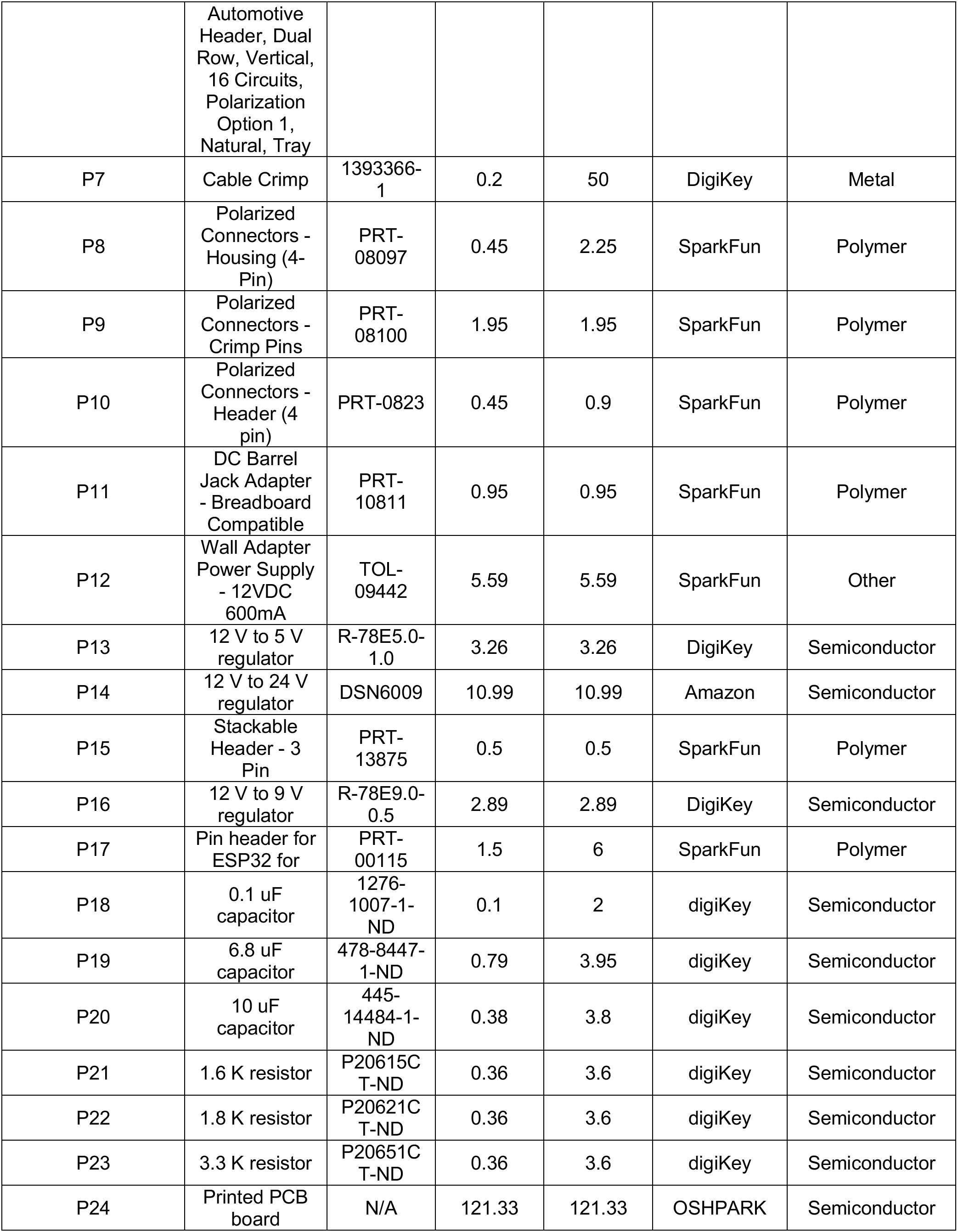
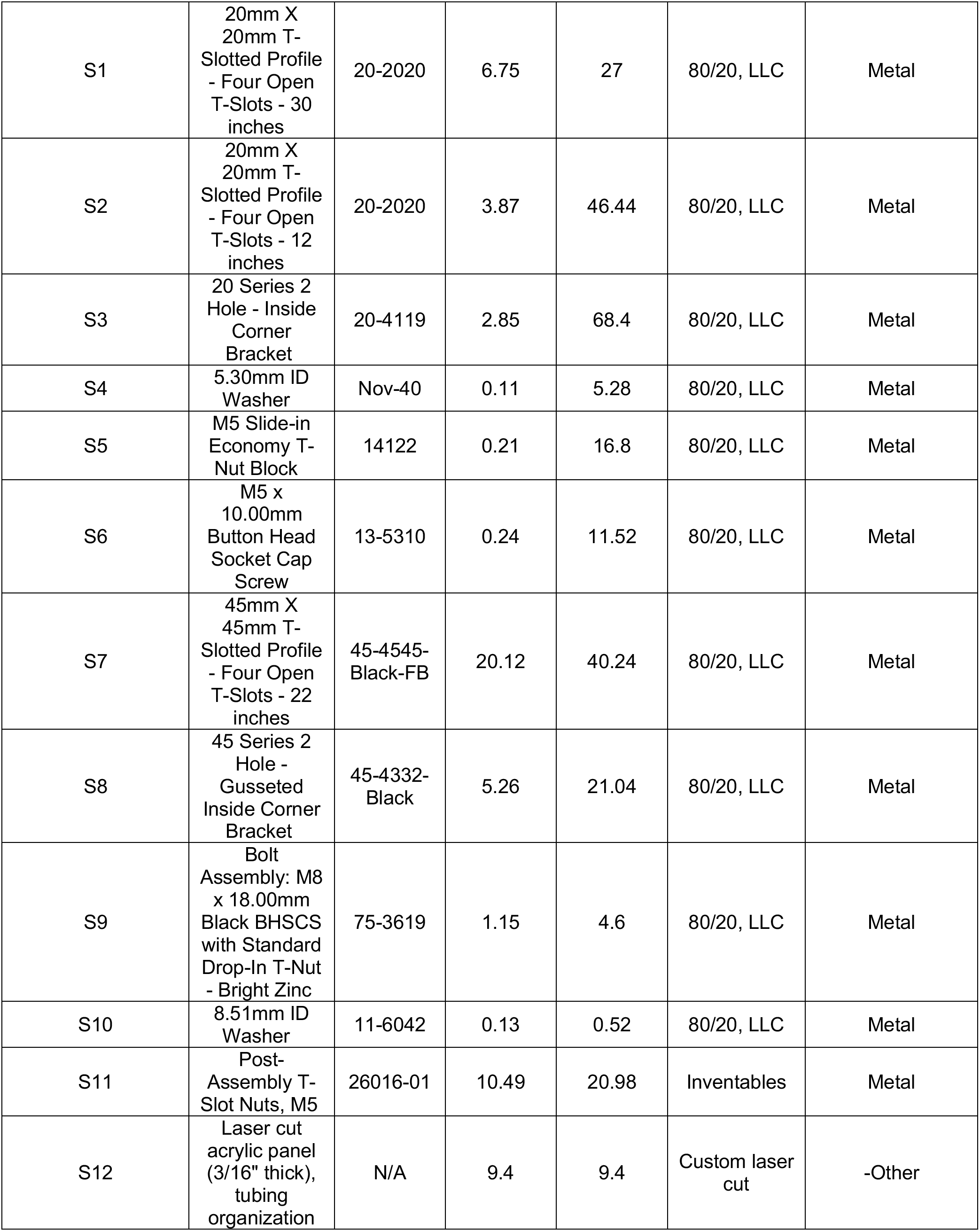
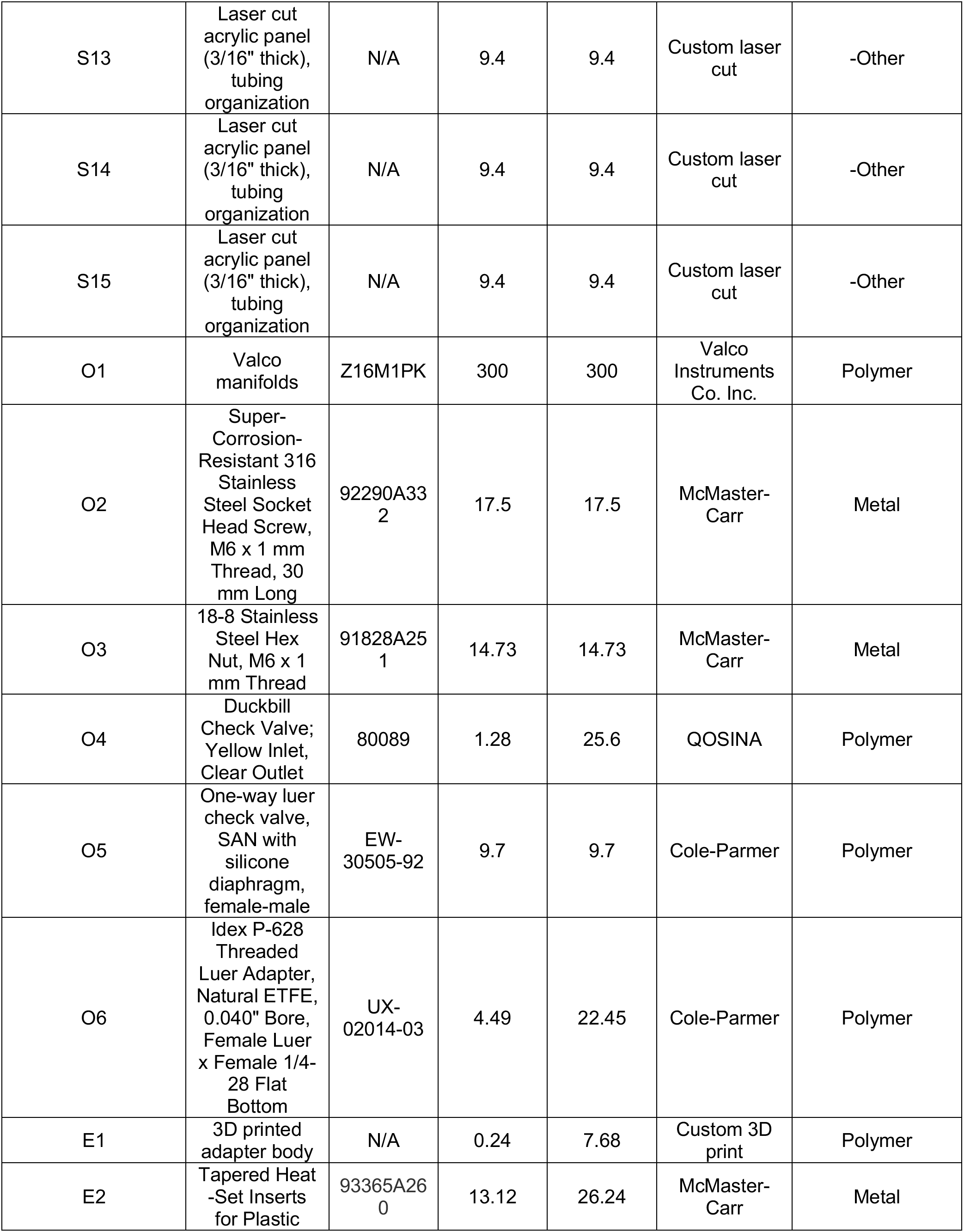
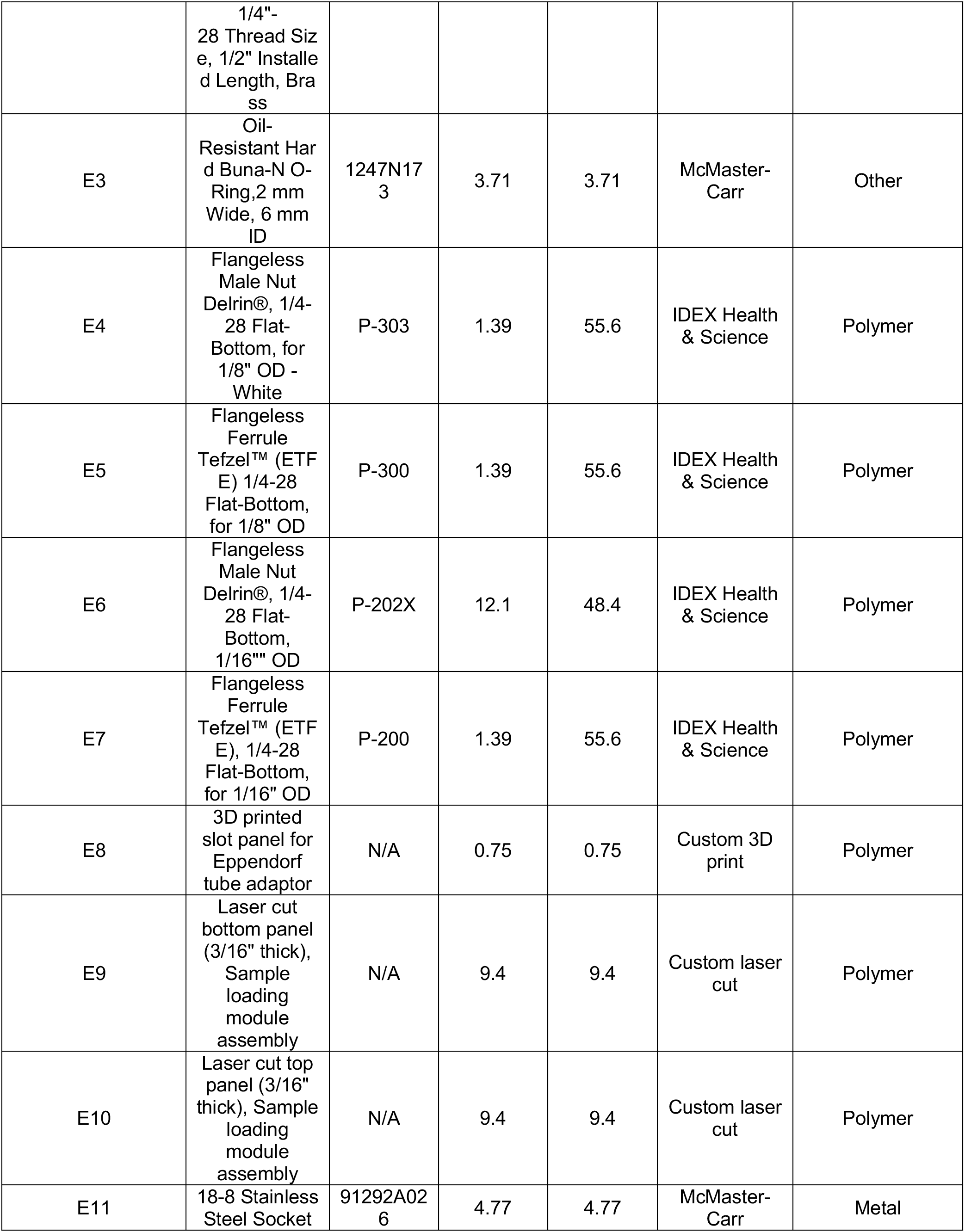
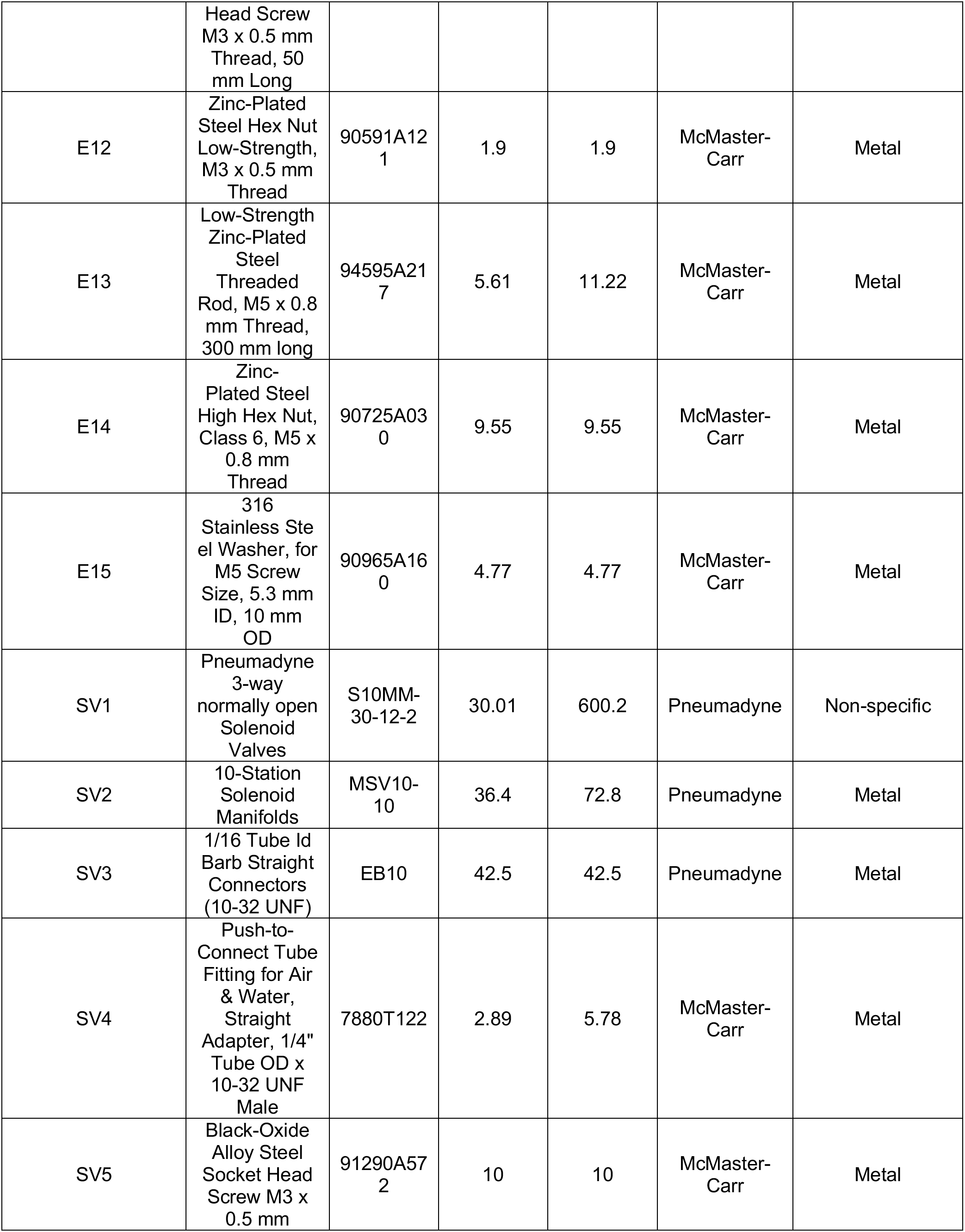
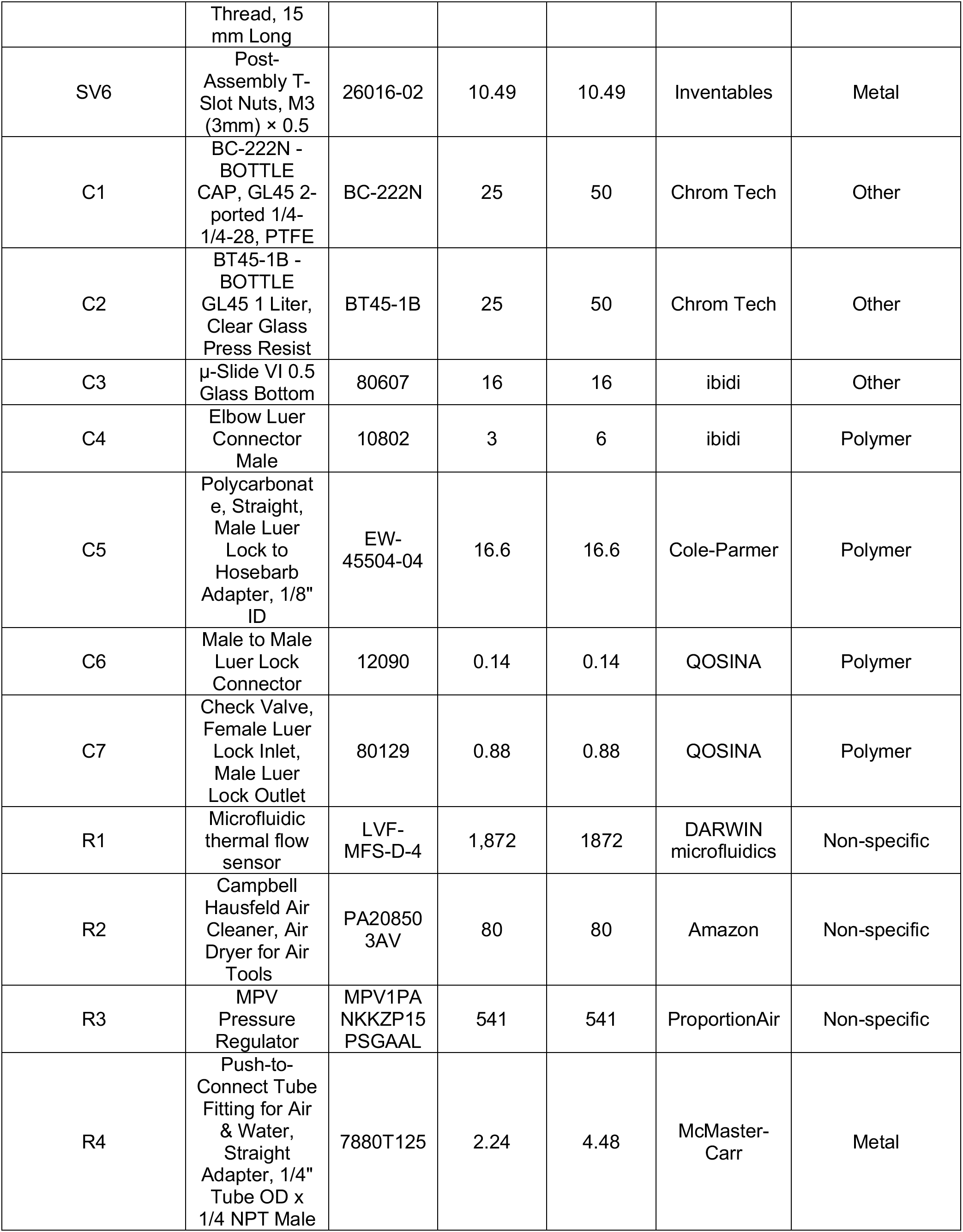
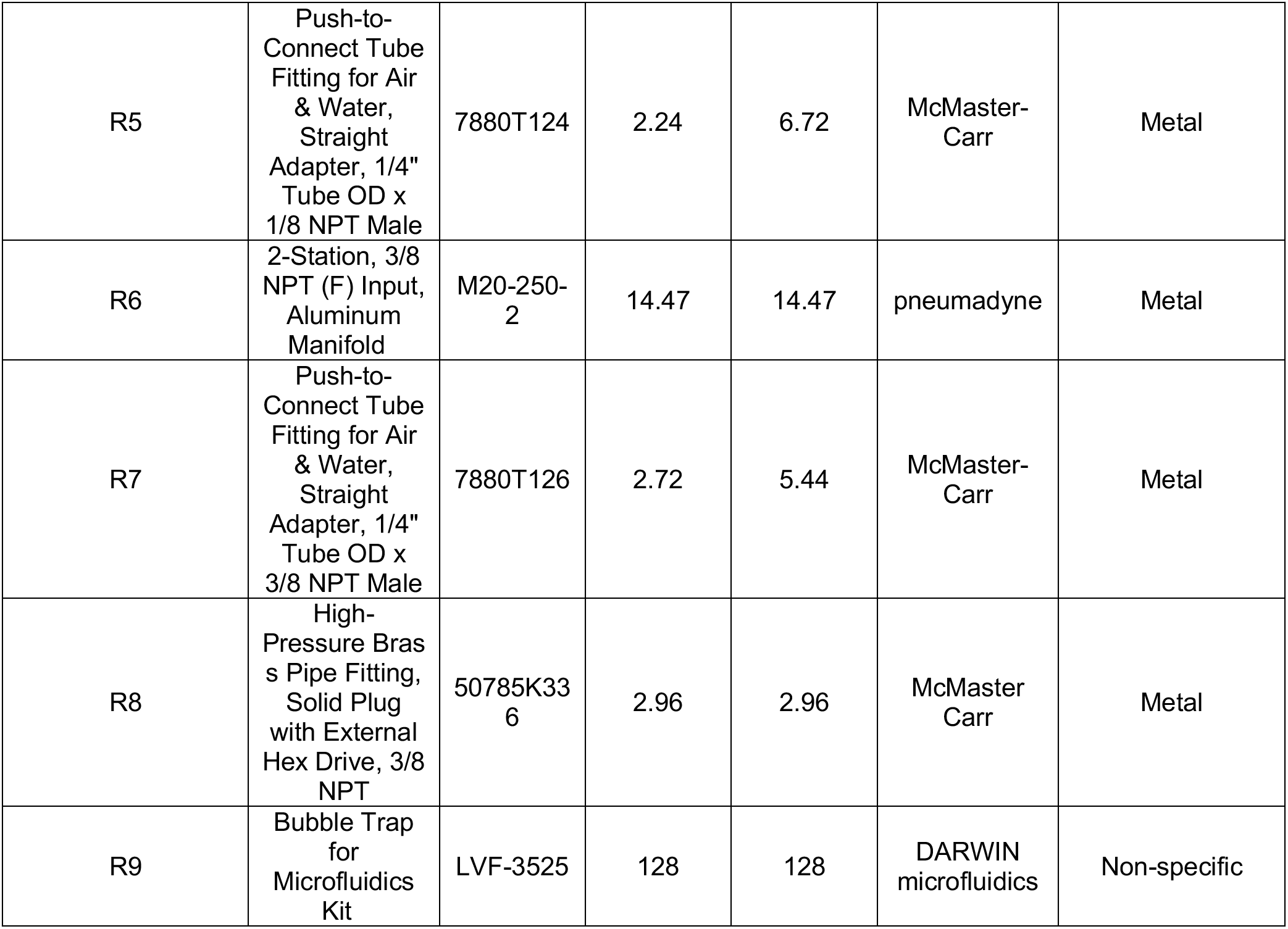

## 5. Build instructions

A step-by-step guide for building the fluidics system is provided in the supplementary materials. There are 7 sections in the guide each describing an individual module and/or assembly with the last section about connecting everything together.

## 6. Operation instructions

A calibration and operation guide is provided in the supplementary materials. With the 14-round fluor oligo exchange to image pan-alpha satellite in Hela cell as example, the instruction describes step-by-step the calibration and operation procedure for setting up the fluidics system for automatic liquid handling for the imaging experiment.

## 7. Validation and characterization

### 7.1 Flow rate control

A robust flow rate control is important for controlling the injection volume. The flow rate is linearly proportional to the amount of pressure that is applied to the reservoir. A calibration routine is integrated into the user interface. This routine can complete the automatic flow rate calibration with the 1.5 ml PBS buffer that is loaded in the 2 mL Eppendorf tube. This helps to minimize the sample loading effort for calibration. By default, 3 target flow rates, 140 μL/min, 630 μL/min (denoted by 600 μL/min in the calibration tab in the user interface), and 995 μL/min (denoted by 1000 μL/min in the calibration tab in the user interface) can be selected from the user interface with tested calibration parameters. The flow rate calibration routine that can calibrate all 16 channels without any human interaction takes about 40 mins. Table 1, Table 2, and Table 3 shows the measured flow rate for all 16 channels for the 3 target flow rates respectively. The average accuracy of the 16 channels is <+/-5% of the target flow rate.

**Table 1.** Flow rate calibration result of target flow rate = 140 μL/min. Mean and standard deviation (SD) are based on 3 measurements.

| Channel no. | Target flow rate = 140 $\mu\text{L}/\text{min}$ | | | |
| --- | --- | --- | --- | --- |
| | Measured flow rate ($\mu\text{L}/\text{min}$) | | Percentage error | |
|  | Mean | SD | Mean | SD |
| 1 | 139.93 | 2.59 | 1.38 | 0.76 |
| 2 | 141.73 | 9.07 | 4.57 | 3.60 |
| 3 | 142.76 | 7.80 | 4.93 | 0.63 |
| 4 | 144.58 | 7.56 | 4.67 | 3.54 |
| 5 | 140.93 | 6.10 | 3.11 | 2.26 |
| 6 | 143.16 | 2.75 | 2.25 | 1.96 |
| 7 | 142.21 | 9.93 | 5.55 | 2.78 |
| 8 | 145.81 | 5.41 | 4.15 | 3.86 |
| 9 | 141.69 | 6.10 | 3.66 | 1.06 |
| 10 | 145.47 | 10.06 | 4.32 | 6.82 |
| 11 | 144.26 | 8.48 | 5.49 | 2.31 |
| 12 | 141.01 | 3.48 | 2.12 | 0.46 |
| 13 | 153.93 | 10.38 | 9.95 | 7.41 |
| 14 | 146.61 | 2.14 | 4.72 | 1.53 |
| 15 | 150.27 | 0.75 | 7.34 | 0.53 |
| 16 | 146.29 | 6.44 | 4.50 | 4.60 |

**Table 2.**
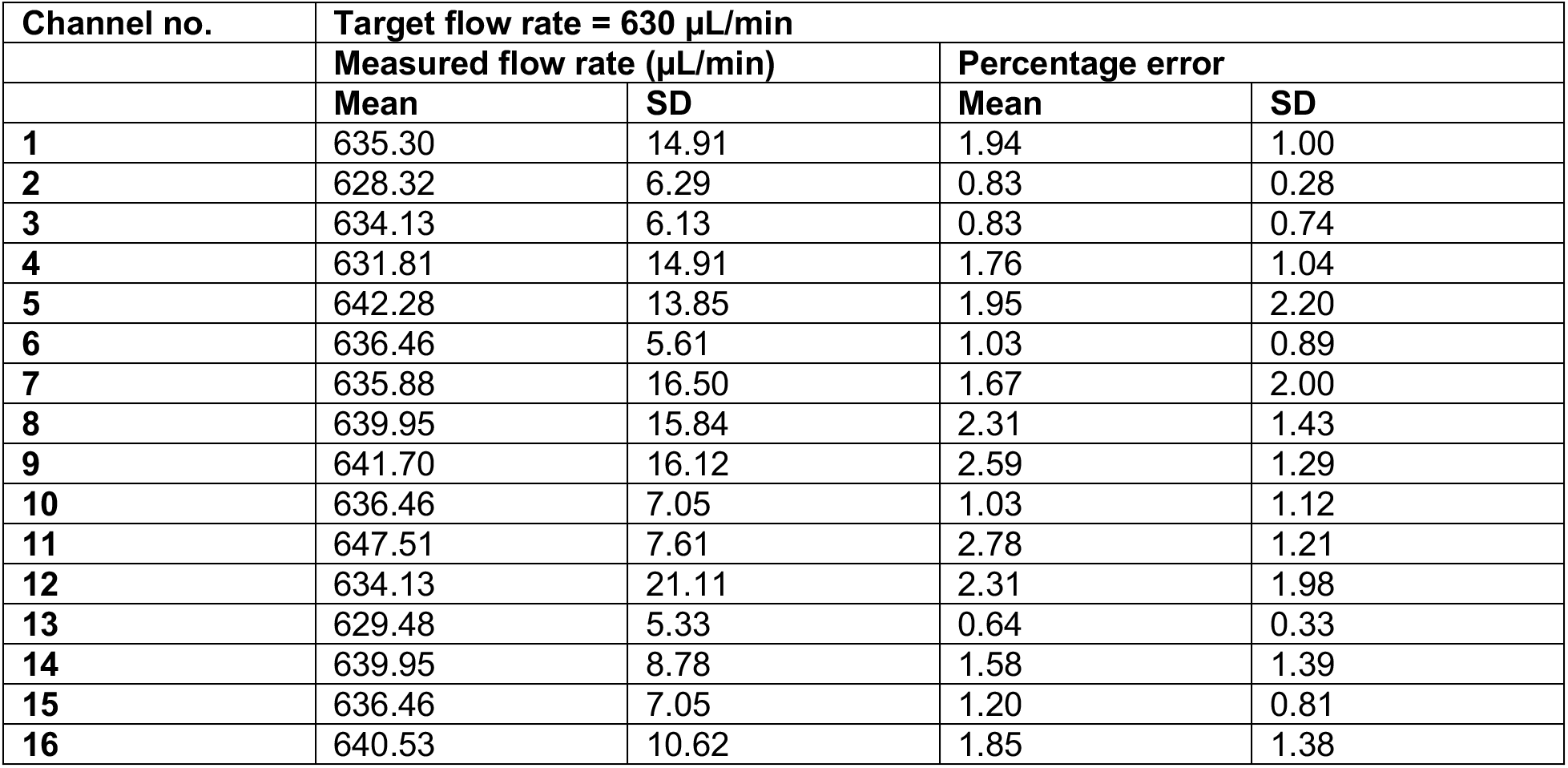
Flow rate calibration result of target flow rate = 630 μL/min. Mean and standard deviation (SD) are based on 3 measurements.

**Table 3.** Flow rate calibration result of target flow rate = 995 μL/min. Mean and standard deviation (SD) are based on 3 measurements.

| Channel no. | Target flow rate = 995 μL/min | | | |
| --- | --- | --- | --- | --- |
| | Measured flow rate (μL/min) | | Percentage error | |
|  | Mean | SD | Mean | SD |
| 1 | 949.47 | 19.00 | 4.58 | 1.91 |
| 2 | 973.90 | 3.63 | 2.12 | 0.36 |
| 3 | 978.09 | 4.19 | 1.70 | 0.42 |
| 4 | 998.34 | 12.27 | 1.05 | 0.21 |
| 5 | 1000.43 | 19.00 | 1.54 | 0.74 |
| 6 | 985.07 | 10.75 | 1.13 | 0.87 |
| 7 | 987.17 | 15.86 | 1.48 | 0.44 |
| 8 | 979.49 | 1.21 | 1.56 | 0.12 |
| 9 | 966.22 | 7.36 | 2.89 | 0.74 |
| 10 | 990.66 | 15.81 | 1.27 | 0.63 |
| 11 | 978.09 | 2.09 | 1.70 | 0.21 |
| 12 | 982.98 | 18.77 | 1.21 | 1.89 |
| 13 | 976.69 | 3.20 | 1.84 | 0.32 |
| 14 | 970.41 | 7.93 | 2.47 | 0.80 |
| 15 | 990.66 | 9.13 | 0.57 | 0.80 |
| 16 | 989.96 | 29.64 | 2.46 | 0.42 |

### 7.2 Minimum injection volume

The fluidic system is designed and built to minimize dead volume (the volume of reagent that used to fill up one individual channel before injecting into the imaging chamber) to reduce reagent cost, especially those of fluorescence oligos. The total dead volume measured by filling one individual channel with DI water is about 200 μL (volume of ~1.5 ft of 0.02” ID tubing (~90 μL) + volume of the one-way check valve (~20 μL) + dead volume of the 16-1 manifold (~1.4 μl) + dead volume of in-line bubble remover (~90 μl)). However, for a channel that is primed with a priming reagent (PBS or DI water), it would be required to inject volume more than this amount to ensure the concentration of the injected reagent when reaching the imaging chamber is maximum (equal to the concentration of the reagent in the Eppendorf tube reservoir). The minimum injection volume (when a channel is primed with PBS) for the reagent concentration to reach maximum in the imaging chamber is empirically determined by an experiment. During the experiment, images of the imaging chamber were recorded every 3 seconds in real-time for each channel from when the reagent (a blue dye is used here, ELVEFLOW Microfluidic Dyes Channels Visualization Kit, SKU: LVF-KXX-06) is injected and the imaging lasted for 75 s for each channel. Between switching channels, 2 mL 1XPBS was injected to wash out the dye from the previous channel and the fluorescence intensity of image of the imaging chamber is verified to decrease to minimum after the washing. The resulting averaged fluorescence intensities change over time for each channel is shown in the plot in Fig. 3. Channel # 1-12, # 15, and # 16 is shown here. Channel # 14 was used for injecting 1X PBS for washing during the experiment. Channel # 13 is not used for this experiment since it is designated for injecting a displacement buffer (for FISH) in the actual sequential FISH experiment. Also, the flow rate is recorded in real-time for estimating the minimum injection volume. The minimum injection volume is estimated as the volume that has been injected when the fluorescence intensity in the imaging chamber first reaches 98% of that in the Eppendorf tube reservoir. Table 4 shows the resulting minimum injection volume for each channel. For all the channels, the injection volume calculated based on this experiment shows to be < 800 μL. Therefore, 800 μL is used as the default minimum injection volume.

**Table 4.** Minimum injection volume for fluorescence intensity reaches plateau in the imaging chamber. This is calculated from Fig. 3. The minimum injection volume is determined by the time at which the fluorescence intensity reaches 98% of the maximum fluorescence intensity in the plot.

| Channel no. | Injection time (s) | Minimum injection volume (μL) |
| --- | --- | --- |
| 1 | 42 | 786 |
| 2 | 42 | 770 |
| 3 | 42 | 793 |
| 4 | 39 | 757 |
| 5 | 42 | 777 |
| 6 | 42 | 774 |
| 7 | 39 | 722 |
| 8 | 42 | 787 |
| 9 | 39 | 703 |
| 10 | 39 | 732 |
| 11 | 42 | 765 |
| 12 | 42 | 760 |
| 15 | 39 | 748 |
| 16 | 39 | 710 |

**Fig. 3.**
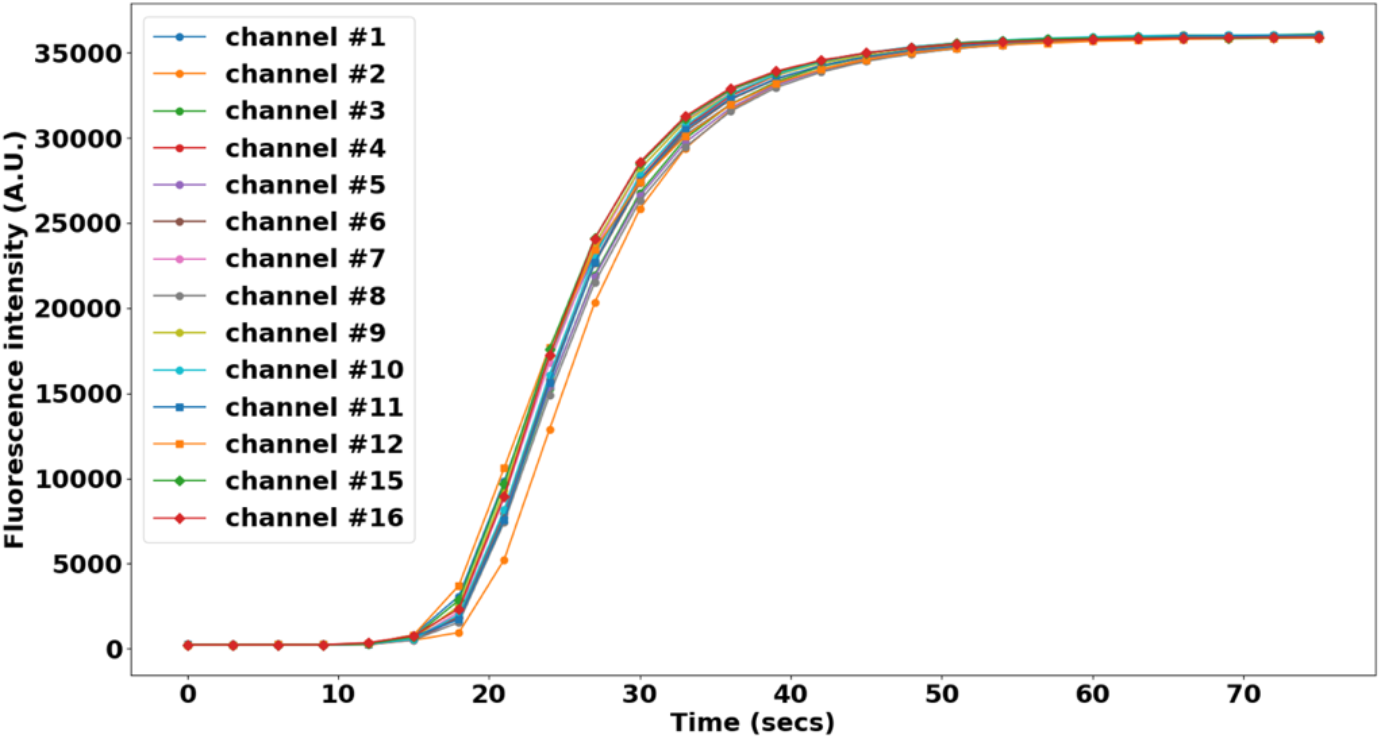
Fluorescence intensity time course during dye injection for 14 channels: channel # 1-12, # 15 and 16. Each data point is the average fluorescence intensity of an image. The images were taken every 3 secs from the onset of the dye injection from one channel till the 75 secs end time. Flow rate is 1 mL/min.

### 7.3 Minimal cross-contamination validation

An experiment was designed and carried out to evaluate the degree of cross-contamination between channels. Reservoirs for channels # 1-12 and 15, 16 were loaded with fluorescent oligos following this order and then repeat it: 488-oligo, 565-oligo, 647-oligo. Reservoirs for channel #13 were loaded with 1X PBS as a washing buffer. And all the channels were primed with PBS. Images were taken in three fluorescence channels (488, 565, 647) after injecting 800 μL (the minimum injection volume from 7.2) fluorescence oligos from one fluidics channel. 2 mL of 1X PBS was injected afterwards and images were then taken again (in three fluorescence channels). These were repeated from the first fluidics channel until the last fluidics channel. Once the reagents are loaded the experiment routine was run automatically with injection commands issued and images taken by NIS-Element. The average fluorescence intensities in the three fluorescence channels are shown in Fig. 4. The order of which fluorescence channel has maximum intensity value follows the order of the fluorescent oligos being injected. Moreover, the fluorescence channels that are not associated with the intended injected fluorescent oligos have very low intensity values. The fluorescence intensity value is also low when 1X PBS was injected to wash the oligo away from the imaging chamber. The fluorescence intensity is very consistent across channels when the same fluorescent oligos is injected. These results show that sequential injection with the fluidics system combined with imaging can be successfully carried out without significant cross-contamination between channels and the concentration of reagent injected from each channel remains consistent.

**Fig. 4.**
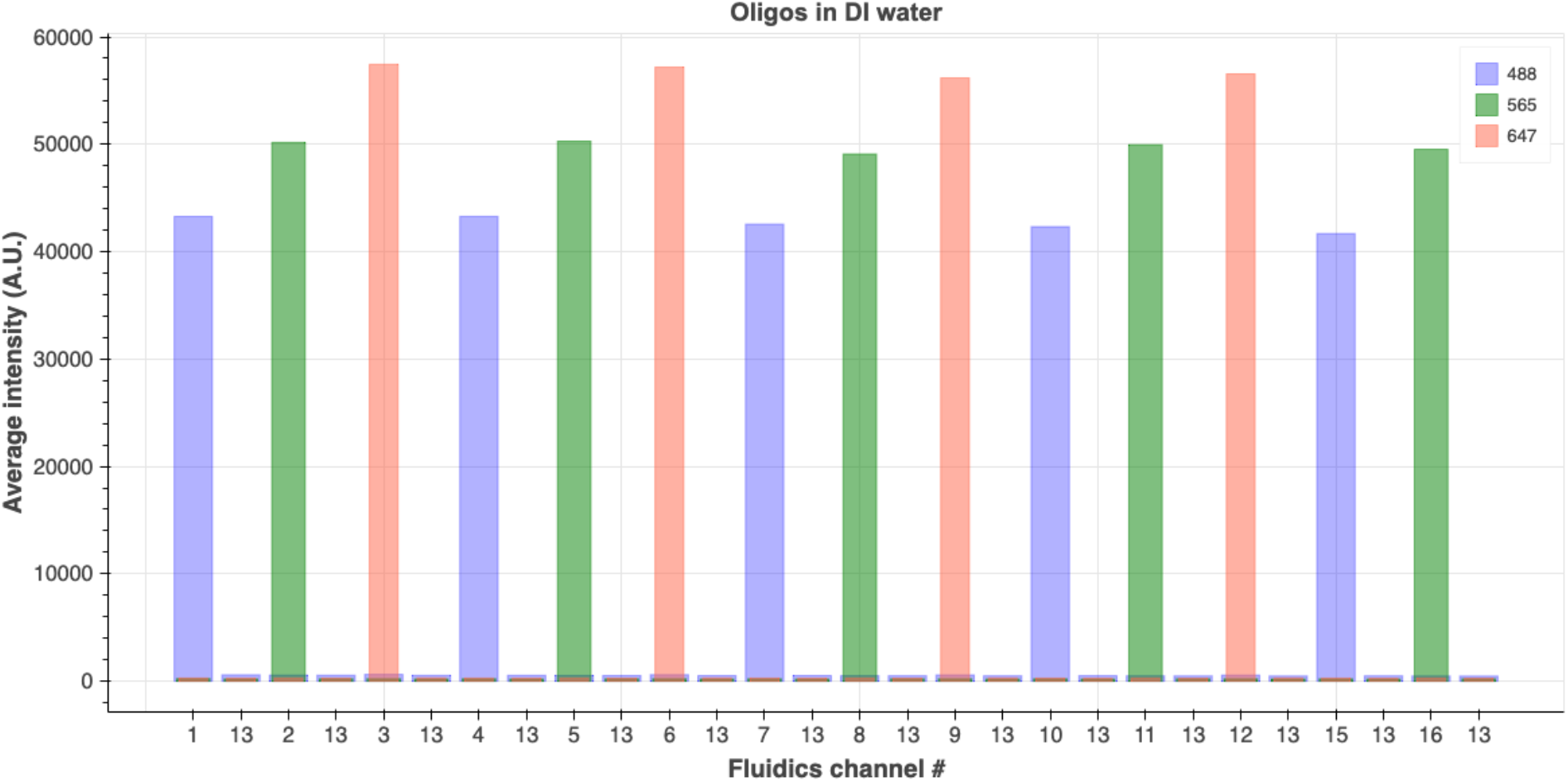
Average fluorescence intensity of images taken in fluorescence imaging channels: 488 nm, 565 nm, and 647 nm. Fluorescence oligos were loaded into the Eppendorf tube reservoirs in this order: 488-oligo for channel 1, 565-oligo for channel 2, 647-oligo for channel 3 and this order was repeated for channels # 4-12 and # 15 and # 16. Images were taken after 800 μL fluorescence oligos reagent from one channel was injected. 2 mL 1X PBS was injected as a washing step to wash out the oligos from the previous injection before injecting fluor oligos from the next channel. Images were also taken after the 2 mL 1X PBS injection finished for that round.

### 7.4 Automating a sequential FISH-based imaging experiment

To demonstrate the fluidics system’s capability of performing oligo probe exchange for sequential FISH-based imaging, an experiment was carried out using DNA-SABER FISH to target the alpha satellite repeat in HeLa cells with three different fluorescent readout oligos. The imaging chamber (1 channel of an ibidi 6-channel slide) was first seeded with HeLa cells and primary FISH probes were hybridized before the imaging chamber was connected to the fluidics system. Each of the primary probes has a SABER [19] concatemer sequence (the ‘p27’ concatemer sequence) that can be targeted by SABER fluorescent oligos as readout probes. Three different SABER fluorescent oligos were loaded into the Eppendorf tubes for fluidics channels # 1-12, channels # 15 and 16 in the following order and then repeat: p27-488, p27-565, p27-647. Reservoirs for channels # 13 and 14 were loaded with a displacement buffer and 1X PBS separately. Fluorescent oligos were injected sequentially and images were taken once the oligos from the current fluidics channel were incubated in the imaging chamber for 1 h (hybridization step) and after the oligos were stripped away by injecting the displacement buffer (washing step). The displacement buffer consisted of 0.4X PBS + 0.04% (vol/vol) Tween-20 + 60% (vol/vol) formamide. The fluorescent oligos exchange and imaging routines were controlled by NIS-Element communicating with the fluidics system’s controller by serial commands. No human intervention was needed once the experiment started to run. A detailed protocol for the fluorescence oligo exchange is included in the supplementary materials. The fluorescence images for the first three rounds of hybridization are shown in Fig. 5 (A). The alpha satellite pattern was illustrated by the corresponding fluorescence oligos. Zoom-in images of one cell from Fig. 5 (A) are shown in Fig. 5 (B). The average pixel intensity was quantified by averaging the fluorescence intensity of pixels belonging to the alpha satellite puncta. This is shown in the plot in Fig. 5 (C). The plot shows that the intensity has the highest value in the imaging channel that is associated with the intended injected fluorescent oligos and the intensity is very low in the other two imaging channels. Also, the intensity from images taken after introducing the displacement buffer is low (images taken after the washing step in each cycle). The decrease of intensity in the plot over cycles is likely due to the instability of the fluorescent oligos in the PBS buffer. Nevertheless, the result demonstrates that the fluidics system is robust in handling complicated fluidics exchange routines for sequential FISH-based imaging.

**Fig. 5.**
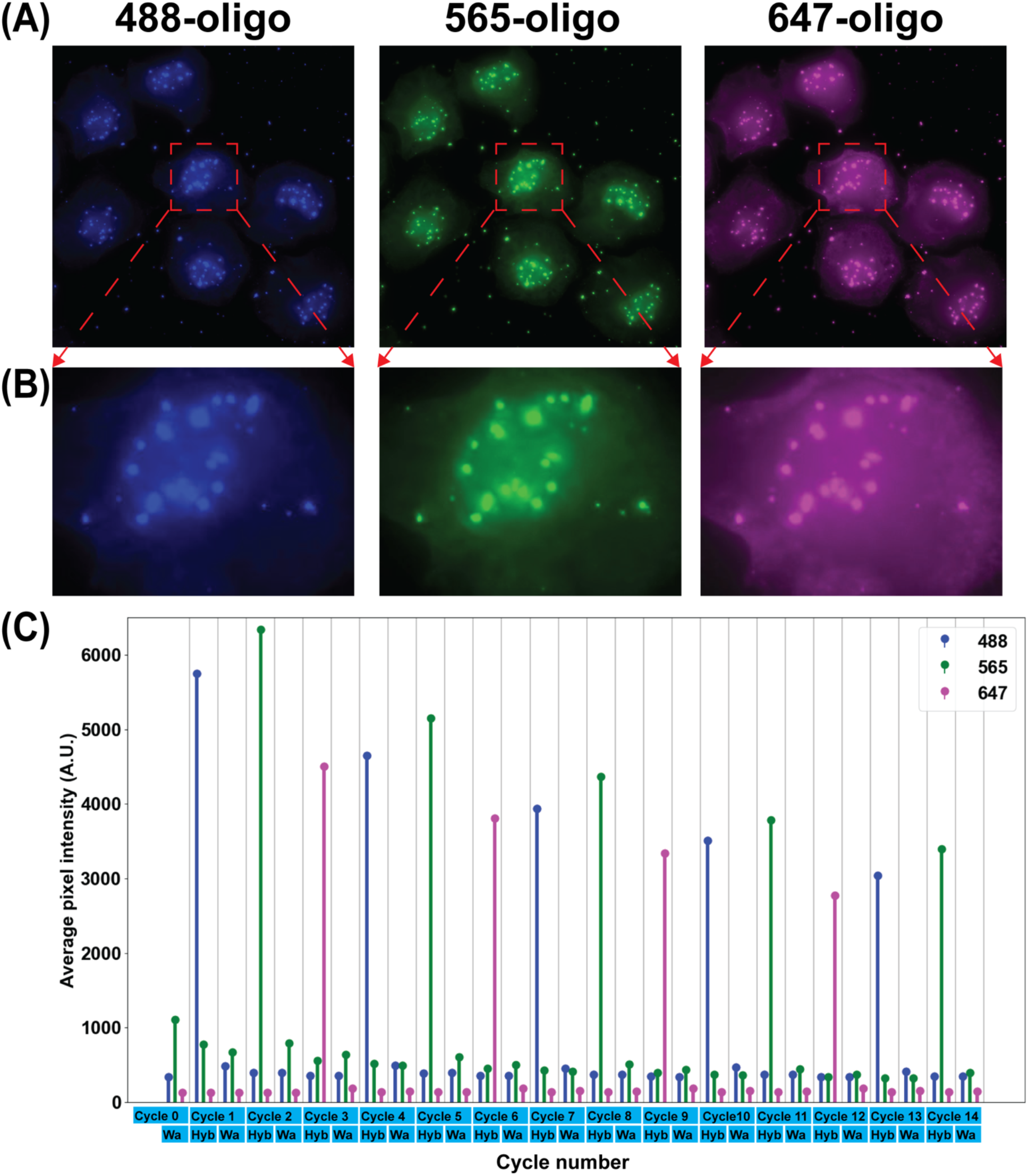
Result of the sequential FISH-based imaging targeting pan-alpha in HeLa cell. (A) Images taken after injecting fluorescence oligos from fluidics channel # 1, 2, and 3 (after the hybridization step in Cycle 1, 2 and 3). (B) Zoom-in images of one cell from (A). (C) The average pixel intensity of pan-alpha puncta in images taken in fluorescence imaging channels 488 nm, 565 nm, and 647 nm in each Cycle. Cycle 1 to 12 inject fluorescence oligos from fluidics channels #1 to 12, Cycle 13 and 14 inject fluorescence oligos from fluidics channels # 15 and 16. Each cycle includes a hybridization step (indicated as Hyb in the x axis) followed by a washing step (indicated as Wa in the x axis) except Cycle 0. Cycle 0 only has a washing step. Images were taken after the hybridization step as well as after the washing step in each cycle.

## Supporting information

Bill of Materials

Calibration and operation guide

Step-by-step building guide

## CRediT author statement

*Zhaojie Deng: Conceptualization, Methodology, Software, Writing-Original draft preparation.* ***Brian J. Beliveau****: Conceptualization, Methodology, Writing-Reviewing and Editing.*

## Acknowledgments

The authors thank Dan Fong (Nikon) for technical support with the Nikon microscope and NIS-Element software, Sahar Attar (Beliveau Lab) for help with primary FISH, Eva Kristine Nichols, Yuzhen Liu, Chris Hsu (Beliveau Lab) for providing ibidi slides with seeded cells, and all the Beliveau Lab members for their helpful feedback on the draft manuscript and figures. We also thank the Maker Space in the University of Washington for providing resources for PCB debugging, laser cutting and 3D printing. This work was supported by a Damon Runyon Dale F. Frey Breakthrough Award (to B.J.B.), the National Institutes of Health (under grant no. 1R35GM137916 to B.J.B.), and a Brotman Baty Catalytic Collaborations Award (to B.J.B.).

## Notes

https://github.com/dzhaojie/Fluidics-system-for-FISH

### Competing Interest Statement

The authors have declared no competing interest.

