## Supplementary material for "An open source 16-channel fluidics system for automating sequential fluorescent *in situ* hybridization (FISH)-based imaging": Calibration and operation guide

**Step-by-step calibration and operation guide**

**Note:**

- In this guide, we run through a step-by-step calibration and operation procedure for an example 14-round sequential FISH-based imaging experiment described in the main text.
- Reference to the sheets named ‘14-channel imaging’ (part names start with capital C), ‘Eppendorf tube adapter array’ (part names start with capital E), ‘Tubings’ (part names start with capital T), and ‘Remaining components’ (part names start with capital R) in the *bill_of_materials.xlsx file* for part names for this guide.
- The codes for the user interface app for automatic flow rate calibration and the NIS-Element Macro codes for integrating Nikon microscope for imaging with liquid handling through the fluidics system can be found in the project’s Mendelay data repository (<https://doi.org/10.17632/tkm4w7wp3v.1>).
- The routines used in this example for fluorescence oligos exchange is adapted from protocol in [1] and are given in the section named ‘**Fluorescence oligo exchange and imaging routines**’ in this guide

**Operation guide:**

Section 1. **System setup**
Step 1. Use new tubings to connect two large reservoirs to channel #13 and #14 with fittings and put them into the 3D-printed holders as shown in **Figure S1**. The part numbers for the bottles and caps are **Part C1** and **Part C2**. The fittings are **Parts E4**, **E5**, **E6** and **E7**. (3D-print holders for the bottles were installed to the base of the structural frame, the design file named ‘large_reservoir_holder.stl’ for the holder can be found in the project’s repository.

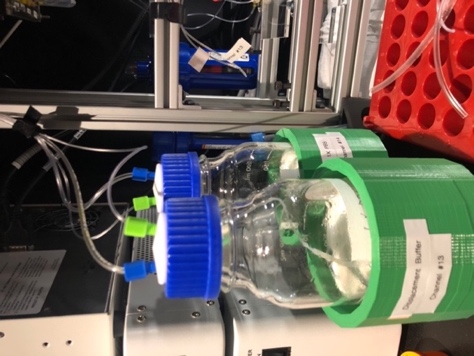

**Figure S1. Large reservoir for PBS and displacement buffer**

Step 2. Use a Q-tip to apply mineral oil to the O-rings of the Eppendorf tube adapters. This is for lubricating the O-ring which makes attaching the Eppendorf tubes to the adapters very easy when loading reagents in the following section.
Step 3. Connect a bubble trap (**Part R9**) between the outlet of the mini manifold (**Part O1**) and the inlet of the flow rate sensor in the flow path.
Step 4. Connect the tubing from the outlet of the flow rate sensor to one end of a channel of an idibi 6-channel slides (**Part C3**) with an Elbow Luer Connector (**Part C4**). With another Elbow Luer Connector (**Part C4**), connect the other end of the channel to a tubing-valve assembly as shown in **Figure S2**. The components in the tubing-valve assembly from top to down are: **Part T4**, **Part C5**, **Part T5**, **Part C6**, **Part C7**, **Part T5**. The one-way-check valve (**Part C7**) is for the purpose of stopping the flow generated due to gravity when all the solenoid valves are closed (when there is no reagent injection happening).

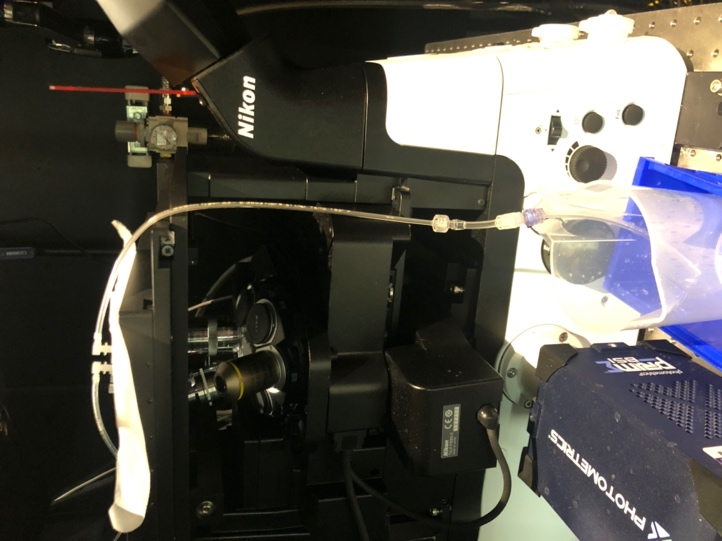

**Figure S2. Waste end connection**

Section 2. **Pre-run priming**
Step 1. Load 14 2 mL Eppendorf tubes each with 1.5 mL 1X PBS onto the fluidics system.
Step 2. Put 200 mL 1X PBS into each of the two large reservoirs. And close the two reservoirs with the caps connected to channel #13 and #14.
Step 3. Connect the main inlet tubing of the fluidics system to a 30 psi pressure source.
Step 4. Connect the fluidics system controller to a computer through the USB port (make sure the driver for ESP32 is installed).
Step 5. Connect the tubing from the outlet of the air filter (**Part R2**) to an air source with pressure of 30 psi.
Step 6. Open the user interface app. Choose a folder to save the calibration data by clicking the ‘Save’ button in the ‘Setup’ tab. Then click the ‘Washing and Priming’ button to start the priming process to prime the fluidics channels with the loaded 1X PBS.

One run of ‘Washing and Priming’ takes about 15 minutes. Two runs of ‘Washing and Priming’ are enough to fully prime the channels and eliminate bubbles from the system.

Step 7. Check the recorded flow rate after each run. The recorded flow rate for all 16 channels will be displayed in the table in the ‘Setup’ tab of the app after the ‘Washing and Priming’ process finished. Each row is the flow rate for one channel measured every 3 secs.

The pressure for ‘Washing and Priming’ is set at 127 (uint8) which is corresponding to 15 psi and the flow rate should be close to the maximum flow rate, 255 (uint8), read out by the flow rate sensor.

Section 3. **Flow rate calibration and verification**
Step 1. Load 14 2 mL Eppendorf tubes each with 1.5 mL 1X PBS onto the fluidics system. (The same set of Eppendorf tubes used for **Section 2** can be re-used in this calibration step)
Step 2. Put 80 mL displacement buffer [1] into a new large reservoir and replace the reservoir connected to channel #13 with it.
Step 3. Inspect flow path, especially the section from the outlet of the mini manifold to the outlet of the ibidi channel slide, to make sure there is no bubble trapped in the channel. If there are bubbles, inject 1XPBS by clicking the ‘Channel_14’ button in the ‘Run single channel’ tab to try flushing out the bubbles. Bubbles tend to trap at the fitting used to connect the tubing to the ibidi channel slide inlet. If bubbles keep sticking at the fitting, try disconnecting the fitting and reconnecting it several times while 1XPBS is being injected until the bubbles are flushed out.
Step 4. Click the ‘Calibration’ tab and select 600 µL/min as the targeted calibration flow rate. Click the ‘Start calibration’ button to start the calibration procedure. The calibration procedure takes about 40 minutes to run. The calibration parameters used here are pre-tested, more information about how to use the app to do calibration for reagents other than PBS and displacement buffer used in this example or other flow rate is presented in **Section 7**.
Step 5. Once the calibration procedure finished. Load 14 2 mL Eppendorf tubes each with 1.5 mL 1X PBS onto the fluidics system. (The same set of Eppendorf tubes used for **Section 2** can be re-used in this calibration step)
Step 6. Click the ‘Calibration verification’ tab and click the ‘Flow rate verification’ button. This step verifies the calibrated flow rate by measuring the flow rate for each channel when applied the calibrated pressure. It takes about 15 minutes.
Step 7. Click the ‘Convert flow rate and show injection time’. The main table in the app window will show the flow rate converted to unit of µL/min for each channel measured in **Step 6** (top row) and injection time (bottom row). Injection volume 800 µL (for fluorescence oligos and displacement buffer) was used to calculate the injection time for this experiment. For 1X PBS, injection volume of 1.6 mL was used to calculate the injection time.
Section 4. **Imaging routine setup (with NIS-Element as example)**
Step 1. Download a .zip file named ‘NIS-Element Macro codes.zip’ from the project’s Mendelay repository (<https://doi.org/10.17632/tkm4w7wp3v.1>) and extract two folders named ‘automatic update Element macros’ and ‘Macros for 16 channel fluidics system V1’. These two folders include custom written NIS-Element macro codes, macro template scripts, a Python script, and a folder with example files generated from flow rate calibration. Two files named ‘WaitTime.npy’ and ‘pr_set_channels_hex.npy’ generated by the user interface app after the calibration and verification from Section 3 are used as input to for the python script ([named ‘update_Element_macro_script.py](XXX)’) for automatically updating the injection time and the set pressure for all 16 channels in the relevant NIS-Element Macro scripts.
Step 2. In the NIS-Element software, open the Macro Panel and set up macro buttons with relevant scripts in the folder ‘Macros for 16 channel fluidics system V1’. Reference to the help document in the actual version of the NIS-Element in use for information about how to set up macro buttons with custom written macro scripts. The Macro Panel used for this example will look like the one in **Figure S3**.

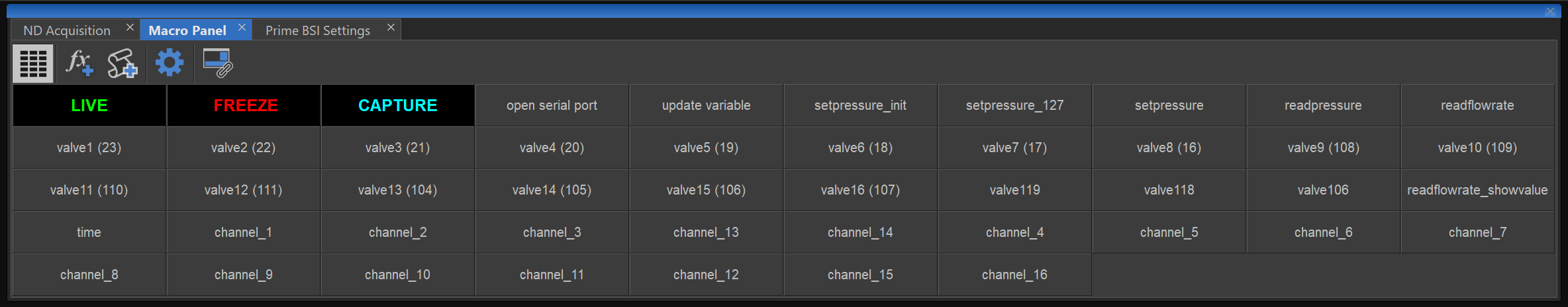

**Figure S3 Macro Panel with macro buttons set up for controlling the fluidics system in the NIS-Element software**

Step 3. Click the ‘Open serial port’ button in the Macro Panel. It might require changing the serial port number (the global variable named ‘*nSerialPort*’) in the macro script named ‘Open serial port.mac’ to the actual serial port number that is assigned to the USB port which the fluidics system is plugged into.
Step 4. Right click the ‘update variable’ button in the Macro Panel to open the macro script associated with this button. The values set for the variables named ‘*WaitTimeIncubation*’, *’WaitTimePBS_sit*’, ’W*aitTimePBS*’, *’WaitTimDB_sit*’, ’*WaitTimeDB*’ are for this specific example experiment. Users can change these values accordingly for their probes exchange handling protocol and then left click the ‘update variable’ button to apply the changes. *’WaitTimePBS_sit*’ and ’*WaitTimePBS*’ work similarly but for different scripts for experiment routines that require two different wait times after injecting PBS. Same with *’WaitTimDB_sit*’ and *’WaitTimDB*’.

Step 5. In this and the next two steps, only settings that are relevant to control the fluidics system are described. Reference to the help document in the software for other settings in the ND Acquisition panel. Open the ND Acquisition panel in NIS-Element, click the Time tab and select it as shown in **Figure S4**. Add 29 phases each with the parameters named ‘Interval ‘set as 2 sec and ‘Loops’ set as 1.

Each phase # is the phase number associated with the time the images were taken in the 3 fluorescence imaging channels. During each phase, serial command was sent from NIS-Element to the fluidics system controller to control which channel to inject. Here phase # 2, 4, 6, 8, 10, 12, 14, 16, 18, 20, 22, 24, 26, 28 are phases when steps (see **Fluorescence oligo exchange and imaging routines** at the end of this guide) designed for fluorescence oligos injection and hybridization were running (hybridization step for **Figure 5** in the main text). Other phases are when steps (see **Fluorescence oligo exchange and imaging routines** at the end of this guide) designed for fluorescence oligos stripping were running (washing step for **Figure 5** in the main text).

Phase #1 here is the washing step for Cycle 0 in **Figure 5** in the main text. Phase # 3, 5, 7, 9, 11, 13, 15, 17, 19, 21, 23, 25, 27, 29 include the washing steps for Cycle 1 to 14 in **Figure 5** in the main text. Phase # 2, 4, 6, 8, 10, 12, 14, 16, 18, 20, 22, 24, 26, 28 include the hybridization steps for Cycle 1 to 14 in **Figure 6** in the main text.

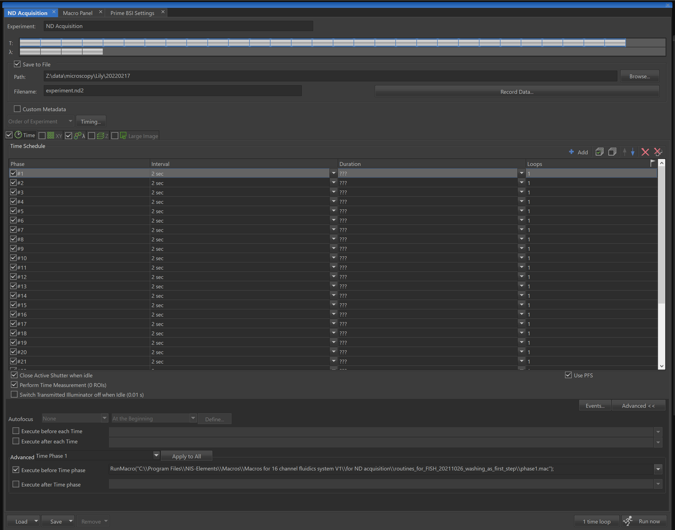

**Figure S4 Setting up time phases in the ND Acquisition panel**

Step 6. In the Advanced section, select ‘Time Phase 1’ from the drop-down menu. Then select ‘Execute before Time phase’. In the text box next to it, put in the command named ‘*RunMacro(“Path to the script”)*’ with “path to the script” pointing to the location of the macro script named ‘[phase1.mac](XXX)’.
Step 7. Repeat **Step 6** for the remaining 28 phases. For example, select macro script named ‘phase2.mac’ for ‘Time phase 2’, select macro script named ‘phase3.mac’ for ‘Time phase 3’, so on and so forth.

**Section 4 Note:** there are 2 issues frequently happening in this version of NIS-Element and below give the work arounds for these 2 issues in order to continue running the experiment smoothly.

1) Sometimes NIS-Element Macro interprets time in second as millisecond. Use the button named ‘time’ in the Macro Panel to test whether the time counting is correct or not. This button runs a script to first set the pressure to maximum, wait for 3 seconds, then set the pressure to zero. Click once this button and observe the macro running status at the bottom of the NIS-Element software. If the time between setting the pressure to maximum and to zero is not 3 seconds, restart the PC and try again. Usually a one-time restart of the PC should fix the issue.
2) Unsuccess serial communication between NIS-Element and the fluidics system controller when the USB of the controller first plugged in. Closing NIS-Element and opening it back up will solve this issue.
Section 5. **Sample loading**
Step 1. Connect an ibidi channel slide with pre-seeded Hela cell with primary probes hybridized to pan alpha satellite (reference to [1] for FISH protocol) to the fluidics system. Use the same method described in **Step 3** of **Section 3** to eliminate bubbles.
Step 2. Load 14 2 mL Eppendorf tubes each with 1.5 mL fluorescence oligos solution (in 1XPBS) onto the fluidics system. For this example, the fluorescence oligos loaded for each channel are listed as in **Table 1***.* Concentration used here is 1 µM.

**Table 1.** Fluorescence oligos loaded for each channel

| Channel # | 1 | 2 | 3 | 4 | 5 | 6 | 7 | 8 | 9 | 10 | 11 | 12 | 15 | 16 |
| --- | --- | --- | --- | --- | --- | --- | --- | --- | --- | --- | --- | --- | --- | --- |
| Fluor oligos | 488 | 565 | 648 | 488 | 565 | 648 | 488 | 565 | 648 | 488 | 565 | 648 | 488 | 565 |

Section 6. **Run experiment**
Step 1. Connect the fluidics system controller to the imaging computer through the USB port.
Step 2. Run ND acquisition in NIS-Element once the cell sample is in focus.
Section 7. **Manual flow rate calibration**

This section describes the procedure to conduct flow rate calibration using the user interface app for reagents other than PBS and displacement buffer used in the demonstrated example (**Section 1** to **Section 6**) and how to extend the calibration routine to achieve flow rates other than the 3 default flow rates.

The calibration routine is designed based on the linear relationship between the applied pressure and the flow rate. There is a coarse calibration followed by a fine calibration. The difference between the coarse calibration and the fine calibration is in the step size of the incremental applied pressure. The coarse calibration determines a range of applied pressure within which a more precise pressure can be determined to generate the target flow rate. The combination of coarse and fine calibrations was designed to minimize the searching time of the required pressure (to achieve the target flow rate) and the amount of the calibration reagent required. The whole calibration routine takes about 40 minutes and can be completed with 2 mL reagent in the Eppendorf tube reservoir.

The pressure search range can be set in the text boxes in the ‘Calibration’ tab. The pressure search range (in the order of start pressure, stop pressure, step size) for channel #1-12 and 16 can be entered in the first row. The pressure search range for channel #15 can be entered in the second row (for this specific built, channel #15 has a much higher flow resistance due to hardware component variability). The pressure search range for channel #13 and 14 can be entered in the third and fourth row (for regular usage in our lab, these two channels are designated for large reservoirs for displacement buffer and PBS). Users can manually change these parameters: start pressure, stop pressure, step size to achieve optimal calibration result. The default values here are verified and tested to be robust for 1XPBS and displacement buffer used in the 14-round sequential FISH-based imaging experiment in the main text.

Users can load the resulting calibration data for individual channel by clicking two buttons named ‘Coarse calibration data’ and ‘Fine calibration data’ and they will be shown in the corresponding tables (first row is the applied pressure; second row is the resulting flow rate). Users can examine these calibration data to identify a more suitable calibration parameter set (start pressure, stop pressure, step size) if the calibrated pressure does not archive the target flow rate. The calibrated pressure for each channel is shown in the table under the button named ‘Calibrated set_pressure’ (In the first row it is shown as Decimal data and in the second row it is shown as Hex data).

As for flow rates other than the 3 default pre-tested flow rates, the python scripts for the interface app provided in the project’s repository can be modified (by changing the value assigned to the variable named FLOW_RATE) easily to adapt the calibration routine to calibrate other targeted flow rates.

**Fluorescence oligo exchange and imaging routines:**

1. Inject fluorescence hybridization solution (fluorescence oligos + 1XPBS) of 800 µl and incubate for 1 hour
2. Wash with 2X 800 µl of 1XPBS
3. Multichannel imaging and take Z-stack
4. Wash with 3X 800 µl displacement buffer: inject 3X 800 µl displacement buffer, then let it sit for 4'
5. Wash with 4X 800 µl of 1XPBS
6. Multichannel imaging and take Z-stack
7. Repeat 1 to 6 for all the fluidics channels.
