## Supplementary material for "An open source 16-channel fluidics system for automating sequential fluorescent *in situ* hybridization (FISH)-based imaging": Step-by-step building guide

**Note:** For convenient reference, all the parts (except parts for solenoid valve array assembly) used in this guide are named with a capital letter and a number. The capital letter for a specific part is the first letter of the name of the sheet in the *bill_of_materials.xlsx* file where the part is listed. For parts for solenoid valve array assembly, their names start with capital letters SV. Most of the parts used to build an assembly/module are listed in a sheet with a name indicating that assembly/module. Tubings are used in multiple assembly/module building sections. Part names for tubings are listed in the sheet named ‘*Tubings*’ in the *bill_of_materials.xlsx* file.

Section 1. **PCB assembly**

Reference to the sheet named ‘*PCB*’ in the *bill_of_materials.xlsx file* for components for this section. Also reference to the PCB schematic file and the PCB layout file in a .zip file (provided in the project’s Mendeley data repository: <https://doi.org/10.17632/tkm4w7wp3v.1>) named ‘*fluidics_system_controller_PCB_design_files.zip*’ for components placement positions on the PCB.

Solder PCB components as shown in **Figure S1**, (A) shows the top side of the PCB and (B) shows the bottom side of the PCB before soldering. Most components are soldered on the top side of the PCB as shown in **Figure S1** (C). An enclosure for the PCB was custom-built by laser cut acrylic panels as shown in **Figure S1** (D). Design files for the enclosure are provided in <https://doi.org/10.17632/tkm4w7wp3v.1>.

Note: the footprint for the ESP32 downloaded from [SnapEDA](https://www.snapeda.com/?gclid=CjwKCAiAyPyQBhB6EiwAFUuakoggojyYhbDRQqJoSp_u3IREyScPOYHsKLifiHcdtkg2V6gFQDfK-RoC4FAQAvD_BwE) (at the time the PCB was designed) was incorrect, fly-wires are used to connect pins on one side of the ESP32 to one of the female header connectors **Figure S1** (C). Make sure the pins from ESP32 are inserted all the way into the headers (this should be the first thing to look at when debugging is needed).

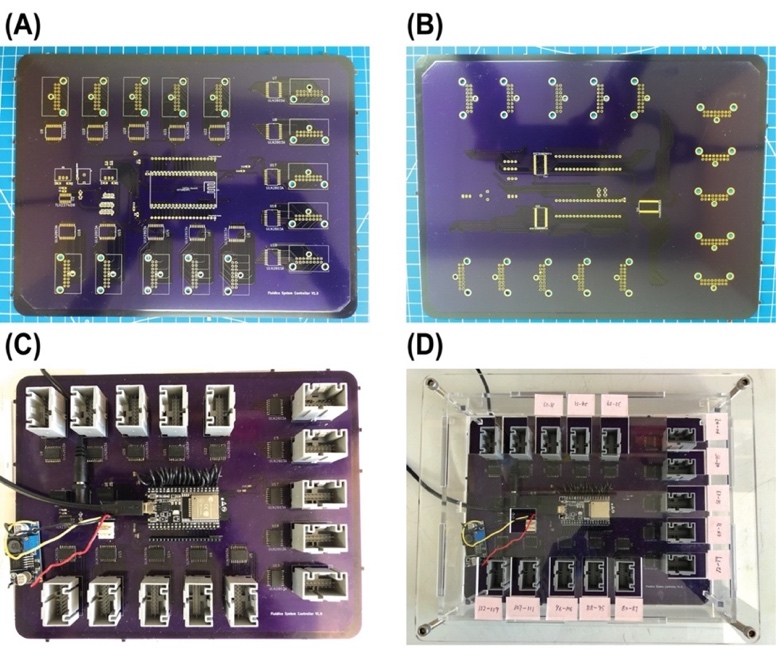

**Figure S1. PCB of the controller before (A and B) and after (C) soldering and secured inside a custom-built enclosure (D)**

Section 2. **Structural frame assembly**

Reference to the sheet named ‘*Structural frame*’ in the *bill_of_materials.xlsx file* for part names for this section. The structural frame refers to the aluminum frame combined with the laser cut acrylic panels that holds all the assemblies/modules and components for the fluidics system for the purpose of tubing organization.

The structural frame consists of 4 vertical positioned 30-inches T-slots (**Part S1**) that define the footprint of the fluidics system. 12 horizontally positioned 12-inches T-slots (**Part S2**) define the 6 levels that hold the major components and assemblies/modules. Except the 3^rd^ level, from bottom to top, the major assemblies/modules and components are: mini manifold, one-way-check valve assembly, Eppendorf tube adapter array, solenoid valve array assembly, controller PCB as shown in **the graphic abstract** in the main text. Between the two 12-inches T-slots at each level (this only applied to the four bottom levels) is one laser cut acrylic panel for tubing organization. The vertical positionings of the Eppendorf tube adapter array, the one-way-check valve assembly, and the mini manifold are critical for proper flow. The Eppendorf tube adapter array is required to be above the outlet of the mini manifold.

Step 1. Begin by assembling the 1^st^ level (the most bottom level) of the aluminum frame. Gather two sets of the parts shown in **Figure S2** (A), **Parts S3, S4, S5, S6** (as in the *bill_of_materials.xlsx file*), another two of **Part S5**, two of **Part S1**, and one of **Part S2.** First, put the two extra **Part S5**s into the slot of **Part S2** as indicated in **Figure S*2*** (B) (The two **Part S5**s here can be replaced post assembly by **Part S11** (**Post-Assembly T-Slot Nuts, M5**), same for other 12-inches T-Slots where two of **Part S5** are required to put into the slot). Then, connect the two 30-inches T-slots (**Part S1**) with the 12-inches T-slot (**Part S2**) by the two sets of parts shown in **Figure S*2*** (A). For the 12-inches T-slot, make sure the side with the two **Part S5**s in it face up. The result for this step will look like **Figure S*2*** (C). Here, **Part S2** is one inch above the bottom of **Part S1.**

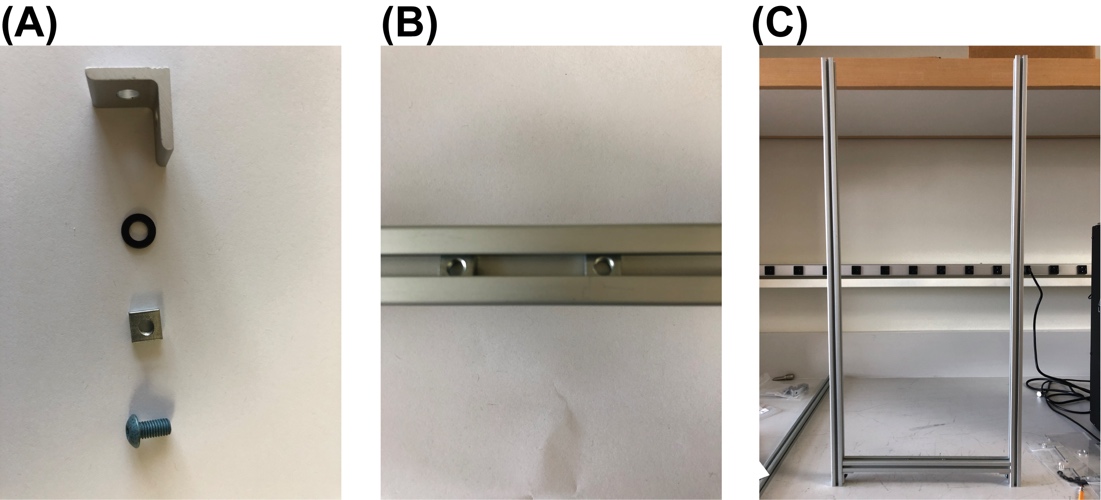

**Figure S2.**  **Structural frame building process 1**

Step 2. Repeat **Step 1** to assemble another side of the frame before moving to the next step. In this step we will get two of the assembled part in **Figure S*2*** (C)
Step 3. Connect the two assembled parts from **Step 2** with one acrylic panel, **Part S12**, as shown in **Figure S3** (A).

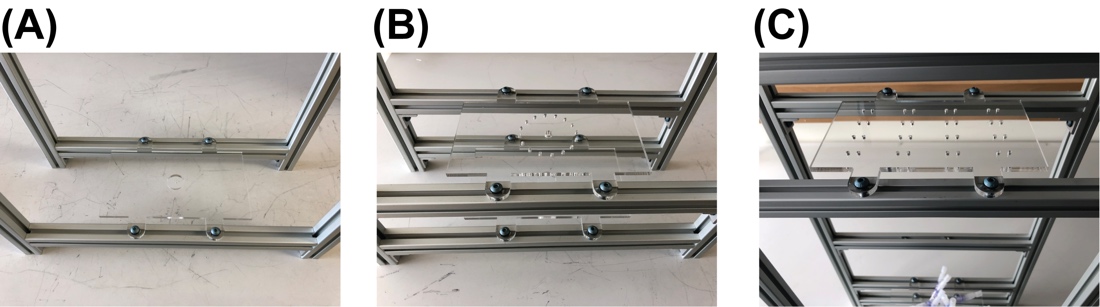

**Figure S3.** **Structural frame building process 2**

Step 4. Continue to install the remaining 12-inches T-slots (**Part S2**) onto the 30-inches T-slots (**Part S1)**. Position each level at a height so that there will be enough space for sample loading and for reaching to all the components when maintenance is required. Remember to put two of **Part S5** into the slot (the one that will be facing up) of each 12-inches T-slot for level 2, 3, 4 (counting from the bottom).
Step 5. Installed the acrylic panel, **Part S13,** to the two 12-inches T-slots on the 2^nd^ level from the bottom as shown in **Figure S3** (B).
Step 6. Installed the acrylic panel, **Part S14,** to the two 12-inches T-slots on the 4^th^ level from the bottom as shown in **Figure S3** (C).
Step 7. This step is optional and should be done after **Section 3** is completed. Installed the acrylic panel, **Part S15,** to the two 12-inches T-slots on the 3^rd^ level from the bottom. The acrylic panel used at this level is for organizing the tubings immediately connected to the one-way-check valves.
Step 8. **Part S7** acts as a base to stabilize the whole fluidics system. Attach one **Part S7** to the bottom end of the structural frame at one side with one set of **Part S4, S5, S6, S8** ,**S9, S10.** Attach another **Part S7** to the other side. This step should be implemented after finishing this building guide if the working area has limited space for accessing all sides of the frame.
Section 3. **One-way-check valve assembly**

Reference to the sheet named ‘*One-way-check valve assembly*’ in the *bill_of_materials.xlsx file* for part names for this section. The one-way-check valves used here are for preventing backflow. They are connected to the inlets of the mini manifold (**Part O1**).

Step 1. Cut tubing (**Part T2**) to length of ~3 inches and insert them into the 16 inlets of the mini manifold (**Part O1**) as shown in **Figure S4** (A).

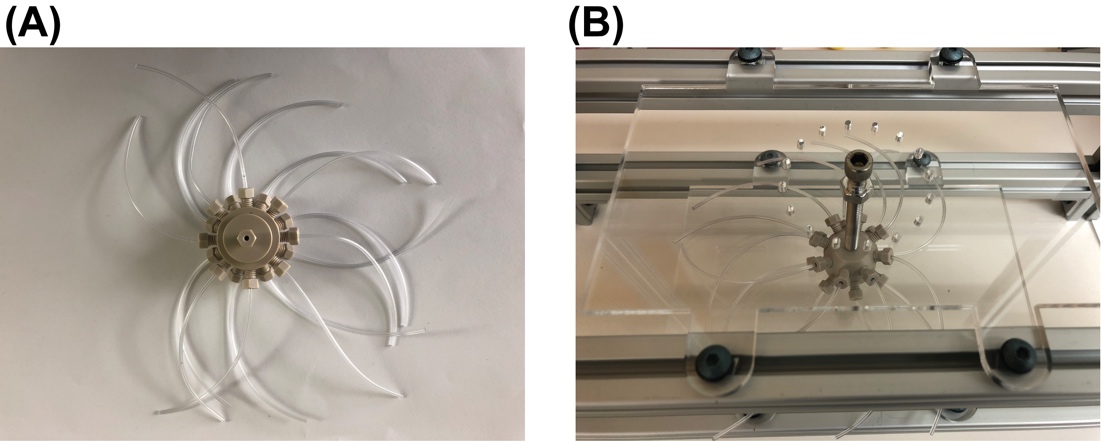

**Figure S4. Install mini manifold on to the structural frame**

Step 2. Install the mini manifold to the structural frame as shown in **Figure S4** (B) with a screw (**Part O2**) and a nut (**Part O3**).
Step 3. Insert the other end of the tubings (that connected to the mini manifold) through the holes in the acrylic panel (**Part S13**)
Step 4. Connect one-way-check valves to the tubings after **Step 3**.

Two different kinds of valve are used (**Part O4 and Part O5**). **Part O4** plus its fittings has a smaller dead volume and is used for channel # 1-12, 15, and 16 for fluorescence probe reagent injection. **Part O5** plus its fittings (two of **Part E6**, **E7**, **O6**, and one of **Part C6**) has a larger dead volume but the fittings provide a more secured connection when applying high flow rate and using buffers with high viscosity like the displacement buffer. Therefore, **Part O5** is used for channel # 13 and 14.

To connect check valve **Part O4**, cut tubing **Part T6** to length of ~3/4 inches, then insert the tubing (that connected to the mini manifold) into **Part T6** as shown in **Figure S5** (A). Then insert both tubings into one end of the check valve as shown in **Figure S5** (B). Here tubing **Part T6** acts as a fitting to secure the connection between tubing **Part T2** and the check valve. There is a direction for check valve **Part O4,** the yellow end is the inlet, the clear end is the outlet

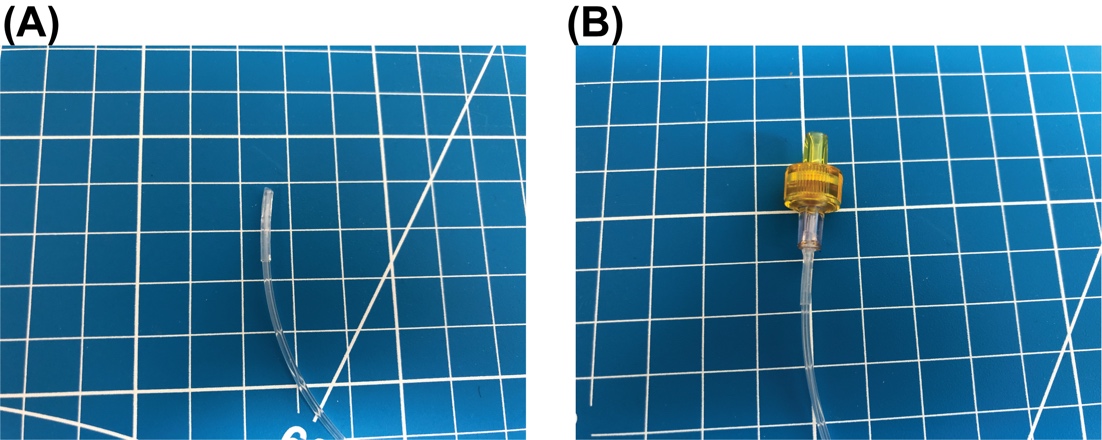

**Figure S5. Connect check valve (Part O4) to the 0.02 “ ID tubing (Part T2) with another tubing (Part T6) cut to length of about 3/4 inches**

Fittings and the check valve **Part O5** are shown in **Figure S6 (A)**. Assemble the parts in **Figure S6** (A) together as shown in **Figure S6** (B). Make two of this assembly. Then connect them to two outlets of the mini manifold.

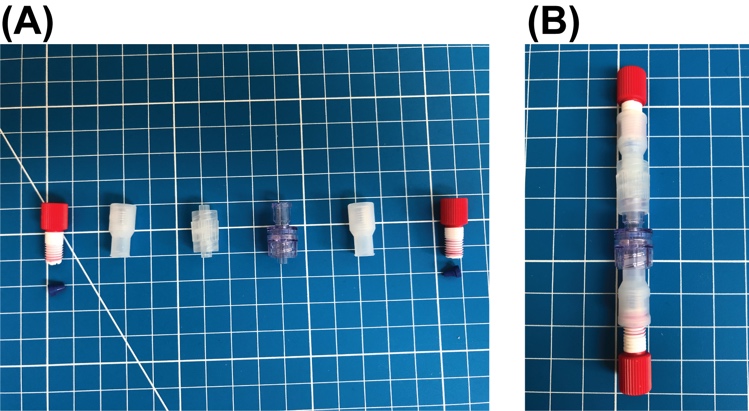

**Figure S6. Connect fittings to check valve Part O5**

**Figure S7** shows the result after all the one-way-check valves are connected to the mini manifold.

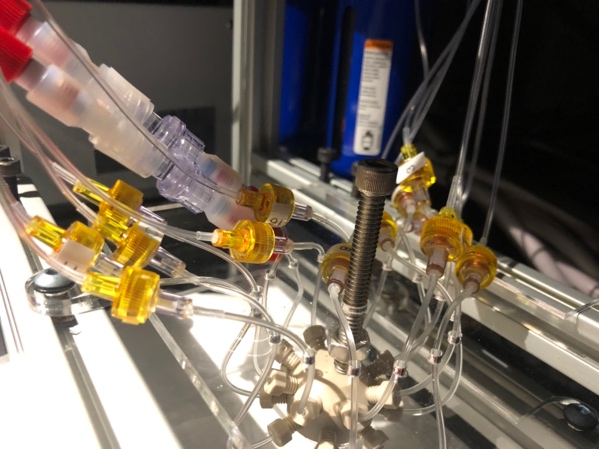

**Figure S7. One-way-check valve assembly**

Section 4. **Eppendorf tube adapter assembly**

Reference to the sheet named ‘*Eppendorf tube adapter array’* in the *bill_of_materials.xlsx* file for part names for this section. The Eppendorf tube adapter is a custom designed and made 3D print part. It enables using 2 mL Eppendorf tubes as reservoirs for positive-pressure-driven liquid sampling application.

Step 1. Install heat-set inserts. Gather one of **Part E1** and two of **Part E2** as shown in **Figure S8** (A). Follow online tutorials (*e.g.*[https://hackaday.com/2019/02/28/threading-3d-printed-parts-how-to-use-heat-set-inserts/)](https://hackaday.com/2019/02/28/threading-3d-printed-parts-how-to-use-heat-set-inserts/) to install the two heat-set inserts **Part E2** into the two holes in **Part E1**. The result for this step is shown in **Figure S8** (B).

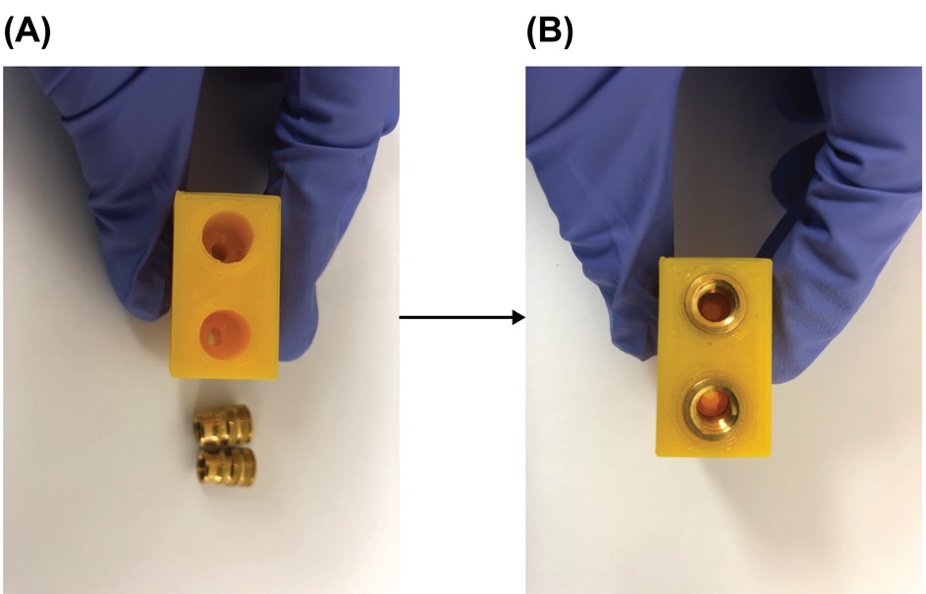

**Figure S8. Install heat-set inserts.**

Step 2. Put on an O-Ring. Gather one of **Part E3** and the resulting part in **Step 1** upside down as shown in **Figure S9** (A). Slide the O-Ring, **Part E3**, onto the extruded port on the adapter body, **Figure S9** (B). Make sure the O-Ring is seated in the designed circular indentation closed to the edge of the port.

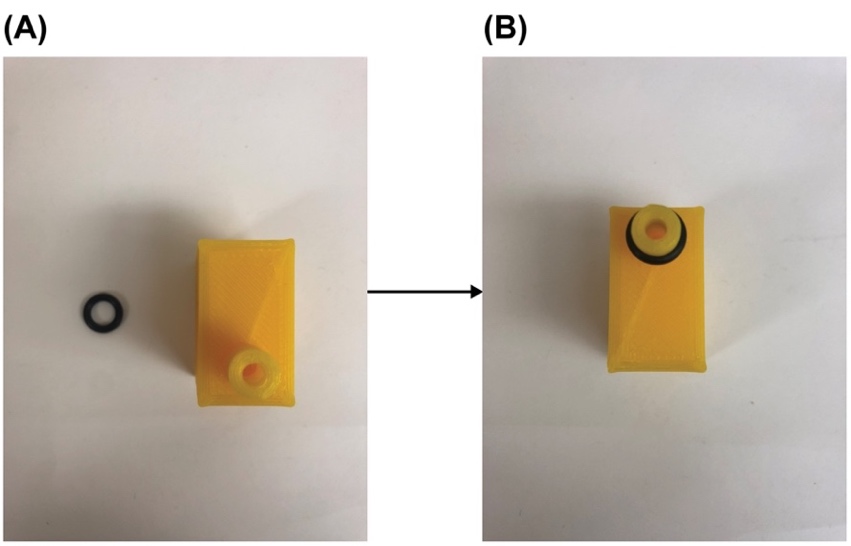

**Figure S9. Put on an O-ring.**

Step 3. Connect the pressure tubing and sampling tubing to the Eppendorf tube adapter. Cut a piece of pressure tubing, **Part T4**, to length of 1 ft. Cut a piece of sampling tubing, **Part T2**, to length of 3 ft. Gather the tubing fittings, **Part E4, E5, E6** and **E7**, one of each. Insert the cut pressure tubing into one of **Part E4** and **E5** as shown in the middle of **Figure S10** (A). Insert the cut sampling tubing into one of **Part E6** and **E7** as shown on the right side of **Figure S10** (A). Use Teflon tape (not shown here) to tightly wrap the thread of the fitting **Part E4** and **E6,** then screw the fittings into the heat-on inserts in the resulting part from **Step 2**. The sampling tubing goes into the hole aligned with the extruded port of the adapter body. And let out around 2 inches length of the sampling tubing from the extruded port. This portion of the tubing will go into an Eppendorf tube to sample reagent. Make sure the tip of this tubing portion just touches the bottom of the Eppendorf tube when the Eppendorf tube is properly hooked up to the port. The result will look like **Figure S10** (B).

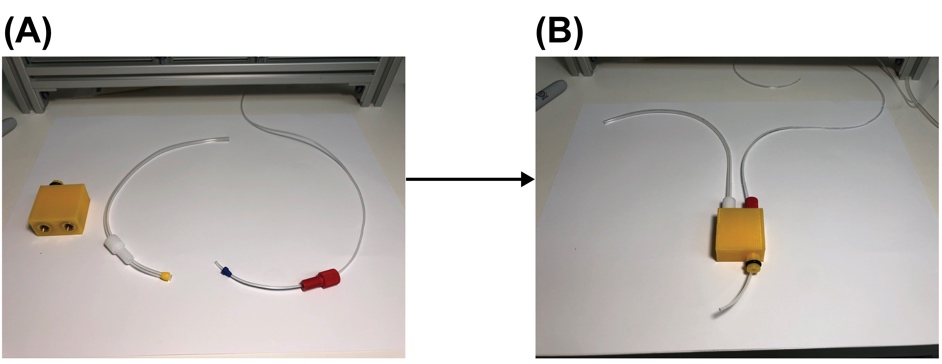

**Figure S10. Connect pressure tubing and sampling tubing to the Eppendorf tube adapter.**

Step 4. Test. Before continuing this step, the firmware should be uploaded to ESP32 of the PCB and make sure commands can be successfully sent to the PCB through USB to control the valves, pressure regulator, and flow rate sensor.

Connect a standard 2 ml Eppendorf tube filled with 1.5 mL DI water or color dye to the adapter as shown in **Figure S11**. Connect the pressure tubing to a pressure source of 15 psi or use the controller and the pressure regulator to regular the pressure to 15 psi. Connect the sampling tubing to the flow rate sensor (**Part R1**) listed in the sheet named ‘*Remaining components*’ in the *bill_of_materials.xlsx* file. Downstream also connects a check valve (**Part C7**) to introduce a flow resistance that is comparable to that used in actual experiment (See **Section 1 Step 3** in the Calibration and operation guide). Use serial commands in a terminal or use the user interface app to read the flow rate through the controller PCB. If the returned flow rate goes over 200 (uint8) when 15 psi pressure is applied then the Eppendorf tube adapter assembly passes the test. If the flow rate is too low, check the seal of the fittings. If you hear loud hissing sounds from the adapter body when applying pressure and the flow rate cannot go over 200(uint8), this means there is a leak in the 3D printed adapter. Printing extra adapters is recommended. Only assemblies that pass the test can be used for the rest of the build.

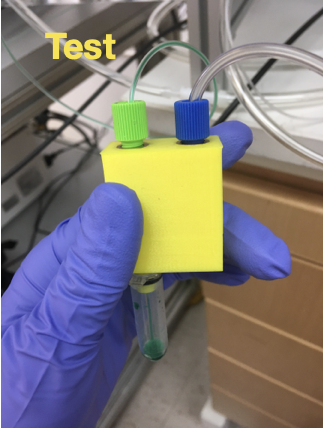

**Figure S11. Test the quality of the Eppendorf tube adapter assembly.**

Step 5. Repeat **Step 1** to 3 to make 20 of the Eppendorf tube adapter assembly. And select 16 of the assembly that passes the test in **Step 4** for the rest of the build.
Section 5. **Sample loading module assembly (Eppendorf tube adapter array)**

Reference to the sheet named ‘*Eppendorf tube adapter array’* in the *bill_of_materials.xlsx* file for part names for this section. The sample loading module assembly combines the 16 Eppendorf tube adapter assemblies into a single module for organized and easy sample loading purposes.

Step 1. Gather **Part E8** and **E9** shown in **Figure S12** (A). Align them as shown in **Figure S12** (B). Flip over both when maintaining the alignment and put them on two supports as shown in **Figure S12** (C). Put one Eppendorf tube adapter assembly built from **Section 4** in this guide into one slot like the one shown in **Figure S12** (C). Finish this step by putting 15 Eppendorf tube adapter assemblies into the remaining 15 slots.

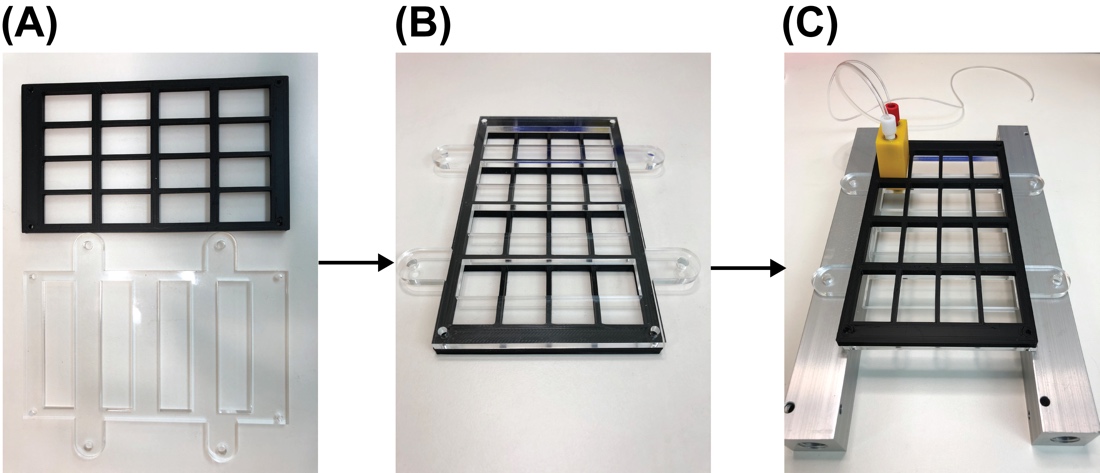

**Figure S12. Align bottom laser cut acrylic panel and the 3D printed slot**

Step 2. Gather **Part E10** shown in **Figure S13** (A) and put it on top of the resulting part from **Step 1** as shown in **Figure S13** (B) with the fittings and tubings coming out of the rectangular holes. Align the screw holes of the top acrylic panel (**Part E10**), the bottom acrylic panel (**Part E9**) with that of the 3D printed slot panel (**Part E8**). Secure the assembly with 4 sets of **Part E11** and **E12** as shown in **Figure S13** (B) and (C).

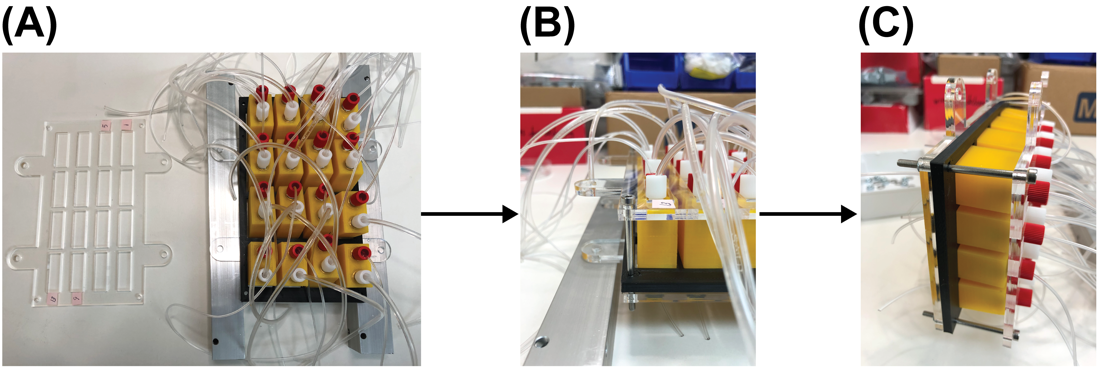

**Figure S13. Install the top laser cut acrylic panel**

Step 3. Cut two of **Part E13 (**a 300 mm long steel threaded rod**)** into half. Gather 16 sets of **Part E14** and **E15.** Insert one of the cut **Part E13** into one of the axillary holes of the acrylic panels and secured with 4 sets of **Part E14** and **E15** as shown in **Figure S14** (A). Then repeat this for the remaining 3 axillary holes of the acrylic panels. Put 2 Post-Assembly T-Slot Nuts (**Part S11**) into the bottom slot of each of the two 12-inches T-slots on the 4^th^ level counting from the bottom of the structural frame. Install the sample loading module onto the structural frame by screwing the 4 long steel threaded rods into those T-Slot Nuts (**Part S11**) as shown in **Figure S14** (B). Insert the tubings into the corresponding holes in the acrylic panel for tubing organization, **Figure S14** (B).

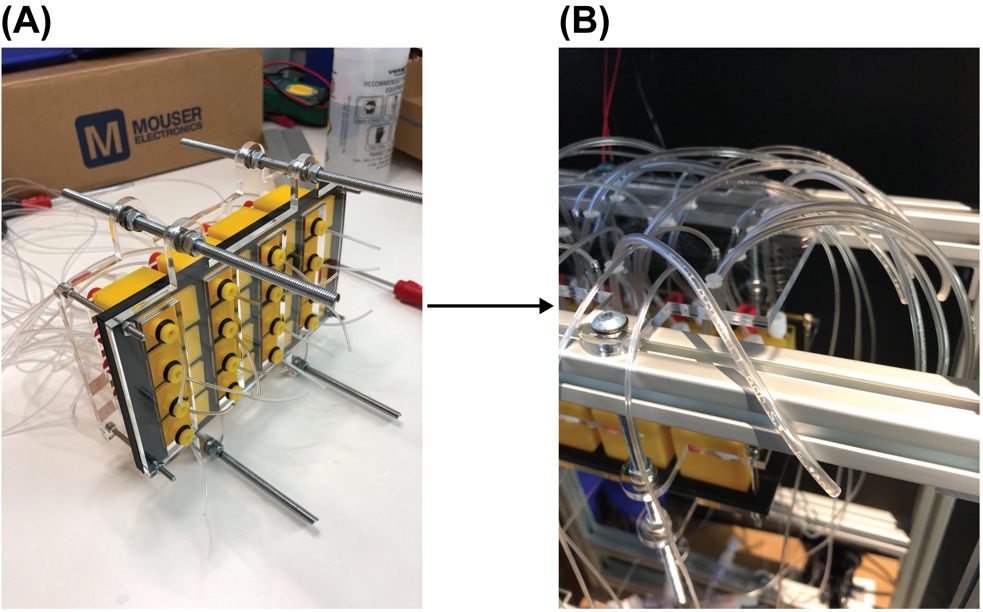

**Figure S14. Install the sample loading module onto the structural frame**

Section 6. **Solenoid valve and manifold assembly**

Reference to the sheet named ‘*Solenoid valve array’* in the *bill_of_materials.xlsx* file for part names for this section.

Step 1. Screw fittings, **Part SV3**, to all 10 output ports of the 10-Station solenoid manifold, **Part SV2.** Install 10 solenoid valves, **Part SV1**, with the screws coming with them onto the manifold as shown in **Figure S15** (A). 4 solenoid valves have been installed in **Figure S15** (A). Though only 8 solenoid valves on one 10-Station solenoid manifold will be in use it is necessary to install all 10 solenoids valves to block leakage through remaining installation holes. Alternatively, 8-Station solenoid manifolds can be used but they were not available at the time when this fluidics system was being built.
Step 2. Screw fittings, **Part SV4**, to the pressure port of the 10-Station solenoid manifold, **Part SV2.**
Step 3. Repeat **Step 1** and 2 to make another solenoid valve array.
Step 4. Put 2 **Part SV6** into the top slot of each of the two 12-inches T-slots on the 5^th^ level (counting from the bottom of the structural frame)
Step 5. Use 2 **Part SV5** to secure each solenoid valve array assembly onto one of the two 12-inches T-slots on the 5^th^ level. Specifically, insert **Part SV5**s into the mounting holes on the manifold (**Part SV2**) and screw them into Post-Assemble T-Slot Nuts (**Part SV6**). The result will look like **Figure S15** (B). Only 8 solenoid valves were installed in **Figure S15** (B) for demonstration but 10 were installed for the finalized build for the reason mentioned in **Step 1**.

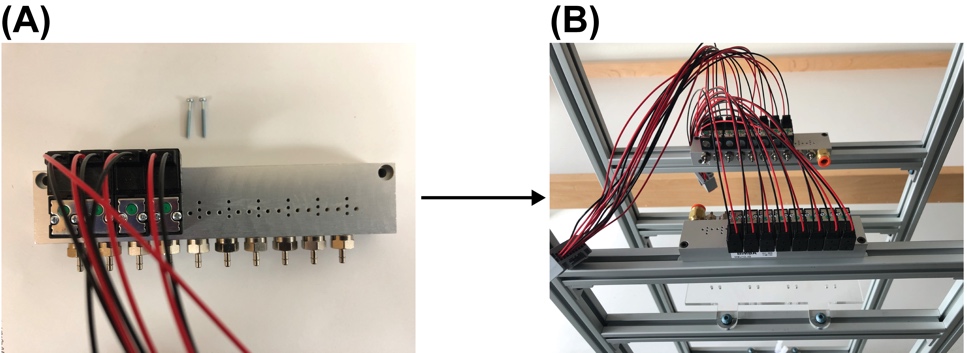

**Figure S15. Solenoid valve array assembly**

Step 6. Cut two pieces of pressure tubing, **Part T4**, each to length of 3 inches. For each solenoid valve array assembly, connect the 2 output ports of the solenoid manifold that are not in use with one of the cut tubings. This is to block leakage through the output ports where the solenoid valves are not in use.
Step 7. Assemble the wires from one solenoid valve array assembly into a connector (**Part P5**).
Step 8. Connect the pressure tubings from the Eppendorf tube adapters to the outlet ports of the solenoid valve manifold and the sampling tubings to the one-way-check valves as shown in **Figure S*16*** (here shows only one-way-check valve **Part O5** for demonstration purpose**,** but in the final built **Part O5** is used only for channel # 13 and 14**, Part O4** is used for the remaining channels as shown in **Figure S7)**

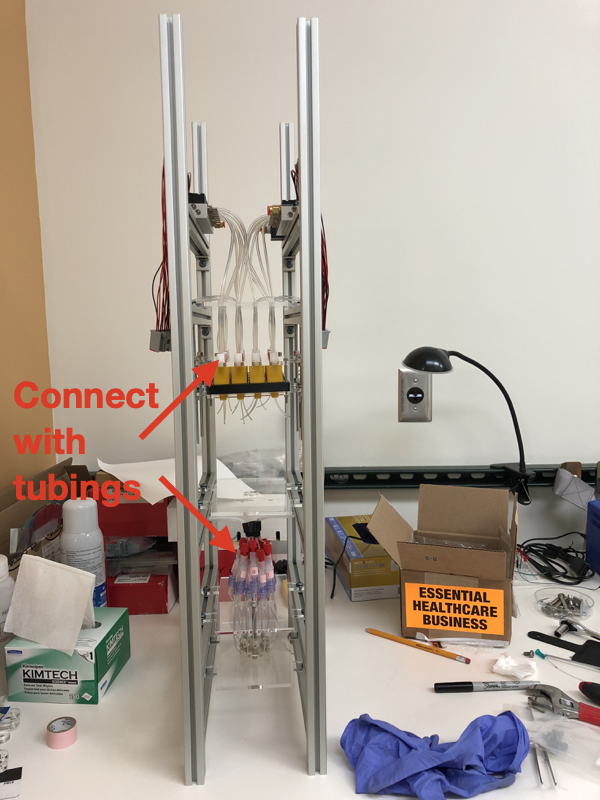

**Figure S16 Connect tubings to solenoid valves and one-way-check valve assembly**

Section 7. **Install remaining components**

Reference to the sheet named ‘*Remaining components’* in the *bill_of_materials.xlsx* file for part names for this section. The mounting screws and T-Slot nuts for installing the remaining components will not be listed here because they are sourced from those listed in previous sections and it should be straightforward to choose the suitable screw and T-slot nut sets to mount/install the remaining components.

Step 1. Install the air filter, **Part R2**, on to the structural frame as shown in **Figure S17** (A). The position of the air filter is required to be closed to the bottom of the whole fluidics system.

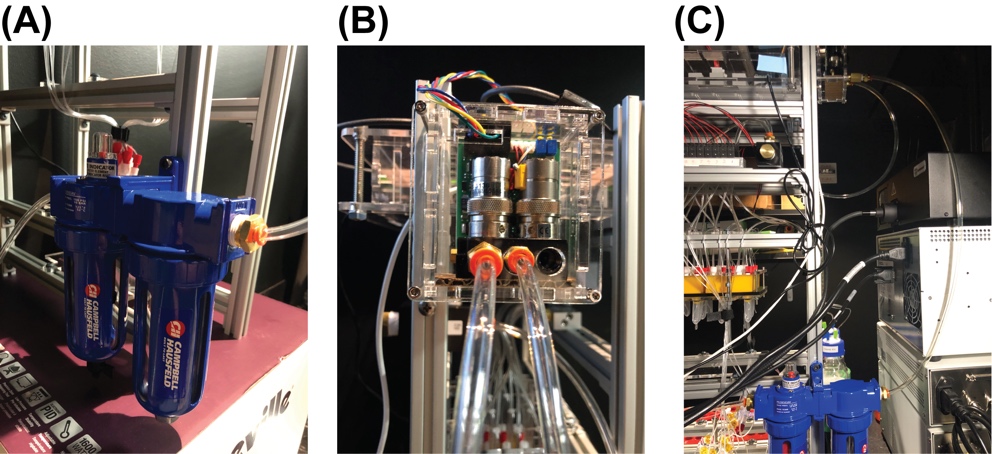

**Figure S17. Install air filter and pressure regulator**

Step 2. Install the pressure regulator, **Part R3**, on to the structural frame as shown in **Figure S17** (B). A custom designed laser-cut case is assembled to house the regulator. The designed file for the case is ‘*case_for_pressure_regulator_laser_cut.pdf*’ in the project repository.
Step 3. Connect the air filter outlet to the inlet of the pressure regulator with tubing, **Part T1** as shown in **Figure S17** (C). The fitting for the inlet and outlet of the air filter is **Part R4**. The fitting for the inlet and outlet of the pressure regulator is **Part R5**. The inlet of the air filter should be connected to a compressed gas source before using the fluidics system for experiment.
Step 4. Connect the outlet of the pressure regulator to the inlet of the splitting manifold (**Part R6**) with tubing, **Part T1**, as shown in **Figure S18** (A). The fitting for the inlet of the splitting manifold is **Part R7**.

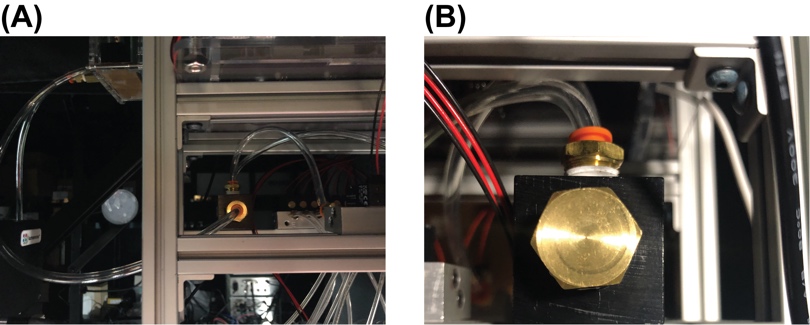

**Figure S18. Connect pressure regulator to splitting manifold**

Step 5. Connect the two outlets of the splitting manifold to the pressure port (**Part SV4**) of the two solenoid valve manifolds with tubing, **Part T1**, as shown in **Figure S18** (A). The fitting for the outlet of the splitting manifold is **Part R4**.
Step 6. Screw the plug, **Part R8**, to the other port of the splitting manifold (**Figure S18** (B)).
Step 7. Mount the PCB controller on the top level of the structural frame using the mounting hole on the case  **Figure S19** (A).

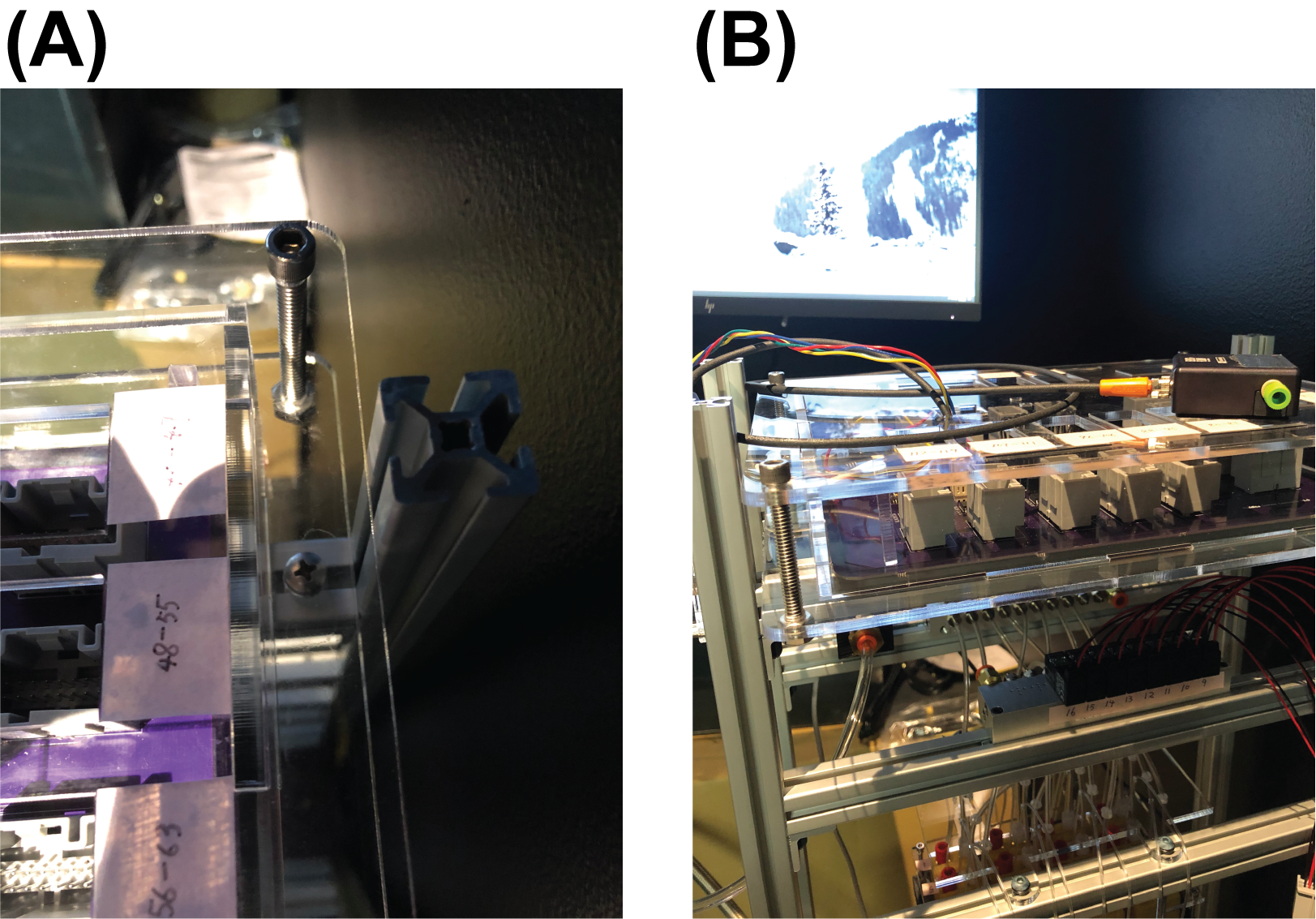

**Figure S19. Mount PCB controller on to the structural frame**

Step 8. Plug the flow rate sensor (**Part R1**) and the pressure regulator (**Part R3**) into their designated 4-pin headers (J6 and J4 in the PCB schematic respectively) on the PCB **Figure S19** (B). The wire for the flow rate sensor must be modified from a M8 connector to be able to plug into the 4-pin header.

A bubble trap (**Part R9**) is recommended to be connected between the mini manifold (**Part O1**) and the flow rate sensor in the flow path.

Step 9. Plug the wire from the two solenoid valve array assemblies into the PCB through their connectors as indicated by the red dash line rectangle in **Figure S20** (A). The PCB has 15 male wire-to-board connectors (**Part P6**) and can control up to 120 solenoid valves. Plug the female wire-to-board connectors (**Part P5**) wired to the solenoid valve array assemblies into any of these male wire-to-board connectors will work as long as the valve number in the NIS-Element Macro code and the user interface app (but **Not** the firmware!!) changed correspondingly. For experiments described in the main text and the operation guide, they are plugged into the 2 connectors indicated by the red dash line rectangles in **Figure S20** (B).

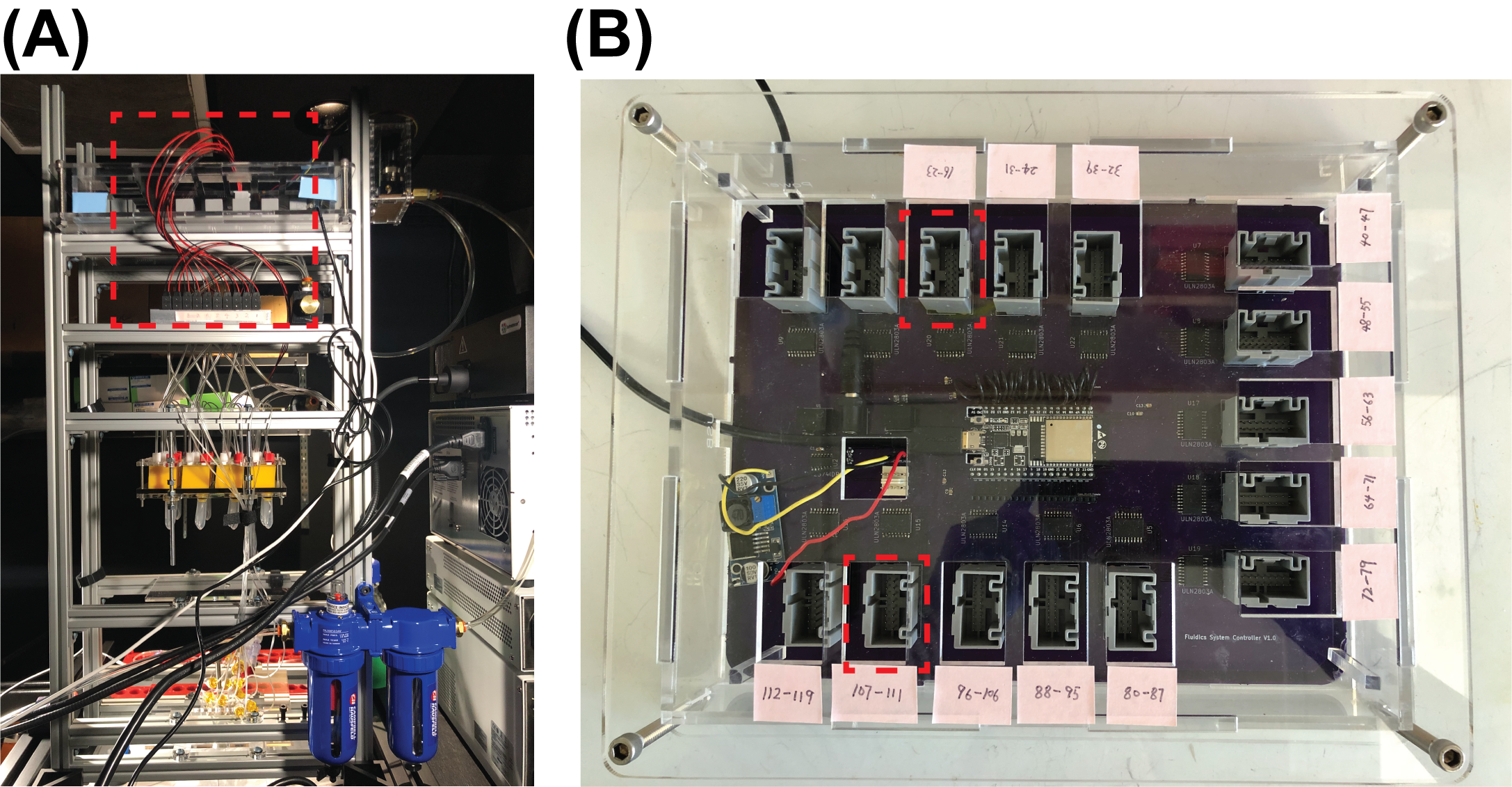

**Figure S20. Connect solenoid valves to the PCB**

Note: The ***ComponentID*** (defined in the firmware code) for valves used here is **16, 17, 18, 19, 20, 21, 22, 23, 104, 105, 106, 107, 108, 109, 110,111**.
